# Mapping glycan-mediated galectin-3 interactions by live cell proximity labeling

**DOI:** 10.1101/2020.05.09.058586

**Authors:** Eugene Joeh, Timothy O’Leary, Weichao Li, Richard Hawkins, Jonathan R. Hung, Christopher G. Parker, Mia L. Huang

## Abstract

Galectin-3 is a glycan-binding protein (GBP) that binds β-galactoside glycan structures to orchestrate a variety of important biological events, including the activation of hepatic stellate cells to cause hepatic fibrosis. While the requisite glycan epitopes needed to bind galectin-3 have long been elucidated, the cellular glycoproteins that bear these glycan signatures remain unknown. Given the importance of the three-dimensional arrangement of glycans in dictating GBP interactions, strategies that allow the identification of GBP receptors in live cells, where the native glycan presentation and glycoprotein expression are preserved, possess significant advantages over static and artificial systems. Here, we describe the integration of a proximity labeling method and quantitative mass spectrometry to map the glycan and glycoprotein interactors for galectin-3 in live hepatic stellate cells. Understanding the identity of the glycoproteins and defining the structures of the glycans required for galectin-3 mediated hepatic stellate cell activation will empower efforts to design and develop selective therapeutics to mitigate hepatic fibrosis.

**Significance:** Because of the weak interactions between individual glycan-binding proteins (GBP), such as galectin-3, and glycans, strategies that allow the direct interrogation of these interactions in living cells remain limited. Thus, the glycan and glycoprotein ligands that are physiologically relevant for galectin-3 binding are insufficiently described. Here, we used a proximity labeling approach that catalytically tags interactors for galectin-3 and identified its pertinent glycan and glycoprotein counter-receptors in live hepatic stellate cells. This study demonstrates that proximity labeling is a powerful tool for mapping GBP complexes in living cells, and when coupled with chemical inhibitors, it can discriminate between protein-protein and protein-glycan interactions.

**Graphical Abstract:** 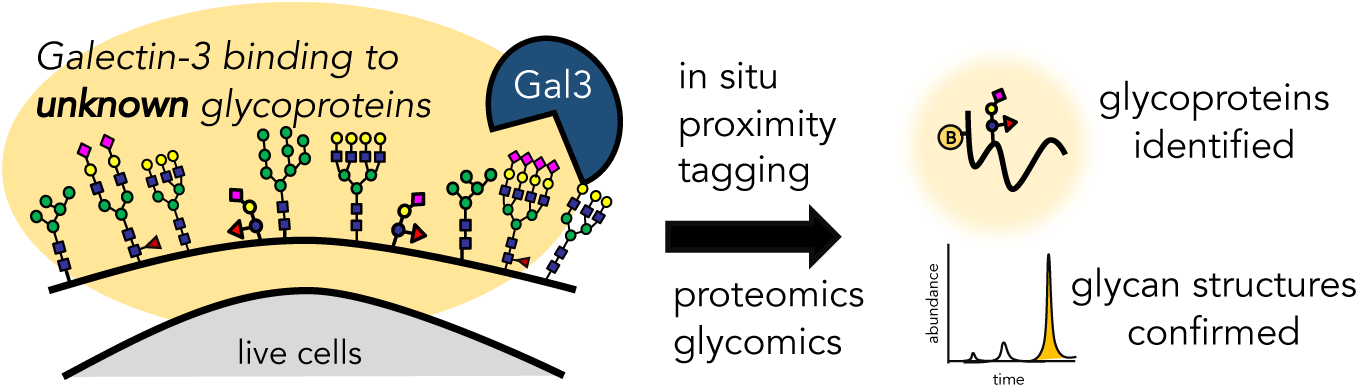

## Introduction

The non-covalent interactions between glycan-binding proteins (GBPs) and glycans dictate many important biological events. Among such GBPs is galectin-3, a β-galactoside GBP that is secreted by macrophages in response to injury [1]. Galectin-3 binds cell surface glycans on hepatic stellate cells (HSCs) and cause their trans-differentiation into muscle-like cells [2, 3], ultimately resulting in hepatic fibrosis, a disease that manifests as the excessive buildup of scar tissue. Despite the critical role of GBP-glycan interactions in this process, the full complement of interacting glycoprotein receptors that are responsible for the fibrotic phenotype remain unknown.

Despite significant advances in glycoscience, the study of GBP-glycan interactions and the identification of glycan-mediated counter receptors remains a recurring challenge. Capturing these binding events often requires some form of artificial reconstitution to amplify individually weak interactions into high avidity binding. Indeed, glycan microarrays with defined mixtures of homogenous glycans or recombinant GBPs have significantly propelled our understanding of glycan-mediated function [4]. Conventional immunoprecipitation and lectin affinity techniques using cell lysates have similarly been used to reveal an initial catalog of 100-185 galectin-3 associated proteins [5-10]. However, these manipulations alter the cell’s native and three-dimensional (3D) configuration and multivalent arrangement, both of which are critically important in the study of GBP-glycan interactions [11, 12].

Another key issue is that of the underlying glycoprotein ligand. Although many glycoproteins carry the glycan epitope for binding a GBP, only a limited set should be recognized as physiologically-relevant receptors, due to the physical constraints imposed by the living cell [13]. While often overlooked, the glycoprotein carrying the glycan can impart specific biological functions to a GBP-glycan binding event [13]. Recent work has put forth the concept of “professional glycoprotein ligands,” where a specific set of glycoproteins (instead of a broadly defined glycome) can exhibit exquisite binding and functional roles [14]. Thus, determining the identity of the underlying core protein that anchors the glycan can be greatly empowering. Not only can it provide an understanding of the 3D arrangement of the glycan (if the three-dimensional structure of the core protein is known), it can also provide additional insight into its expression levels in different cell types and tissues, further informing strategies for selective drug development.

Thus, comprehensive approaches strategies that permit the study of GBP-glycan interactions in live cells while simultaneously facilitating the identification of the physiological glycoprotein receptors possess great potential to impact glycoscience. We hypothesize that proximity labeling strategies [15] using an engineered ascorbate peroxidase, APEX2 [16], could be compatible for elucidating glycan-mediated GBP-glycoprotein interactions. In this approach (**Fig. 1**), APEX2 is fused to a protein of interest, followed by the treatment of cells with biotin-phenol and subsequently with hydrogen peroxide (H_2_O_2_). Under these conditions, APEX2 catalyzes the formation of highly-reactive, short-lived (< 1 ms) and proximally-restricted (< 20 nm) biotin-phenoxyl radicals that covalently tag nearby electron-rich residues. The biotinylated proteins can then be enriched and identified using quantitative mass spectrometry (MS)-based proteomics. Because the (glyco)proteins adjacent to the APEX2 fusion protein are preferentially biotinylated, the resulting MS data provides a readout of its immediate environment. We reasoned that proximity labeling could offer significant advantages over other approaches for determining GBP-glycan interactions, including the opportunity to perform the labeling in live cells and the ability to tag weakly bound glycan-mediated interactors, as the covalent biotinylation reaction ensures that the enrichment step no longer relies on weak GBP-glycan interactions alone. When coupled with inhibitors, the proximity labeling strategy can also distinguish between glycan-mediated and non-glycan mediated interactors. Integration of this approach with quantitative mass spectrometry (MS)-based proteomics would also expedite the assignment of the interacting proteins and aid in providing calculable measures to distinguish interactors from nonspecific binders.

**Figure 1.**
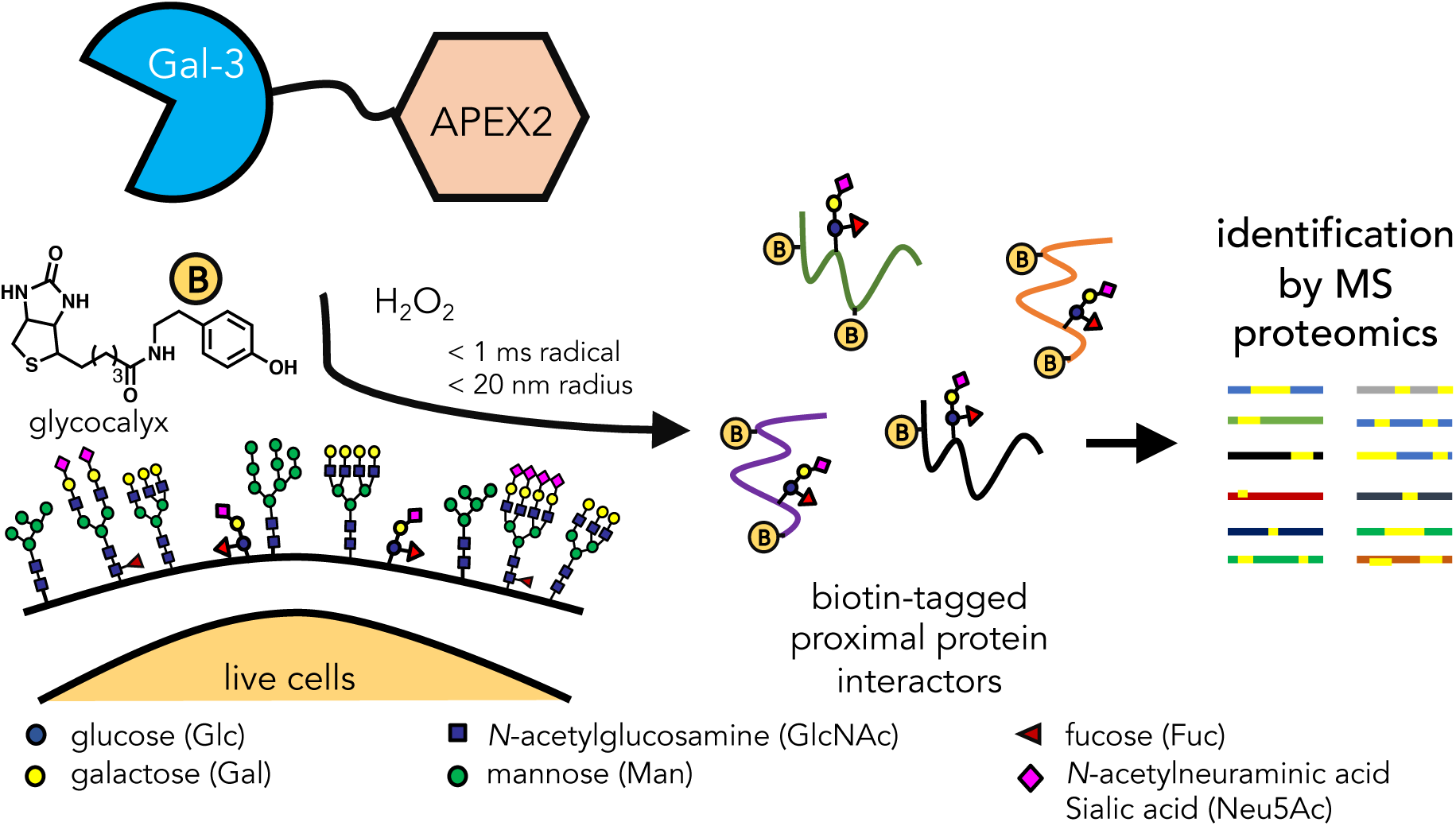
Schematic illustration of the identification of galectin-3 (Gal-3) interacting proteins by in situ proximity labeling. Recombinant APEX2 and galectin-3 fusion proteins are applied to living cells where galectin-3 can freely diffuse to bind its cognate ligands. Upon addition of biotin phenol (yellow circle with “B” symbol; 30 min) and hydrogen peroxide (H2O2; 1 min), APEX2 catalyzes the formation of highly-reactive biotin-phenoxyl radicals that react within a short range (< 20 nm) of the galectin-3 complex within a short time frame (< 1 ms). The biotin-tagged protein interactors can then be identified using mass spectrometry (MS)-based proteomics.

Here, we demonstrate that an APEX2 and galectin-3 fusion protein (PX-Gal3) provides a sensitive and comprehensive approach to map the proteome-wide glycan-mediated galectin-3 interactome in live HSCs. We determine that the exogenous incubation of cells with PX-Gal3 led to glycan-dependent interactions, whereas its cellular over-expression does not. We further validate the interactions between galectin-3 and candidate proteins in vitro and discover that some proteins are direct glycan-mediated receptors. Using MS-based glycomics, we also examined the glycomes of HSC surfaces, PX-Gal3 tagged glycoproteins, and an individual glycoprotein receptor for galectin-3. These results highlight the utility of the in situ proximity labeling approach to discover physiologically-relevant GBP interactors in living cells.

## Results

### Proximity tagging to identify galectin-3 interactors in hepatic stellate cells

We genetically fused the peroxidase APEX2 enzyme [16, 17] to either full-length galectin-3 (PX-Gal3) or a mutant galectin-3 that lacks the N-terminal domain (PX-Gal3_Δ116_) (**Fig. 2A**). This mutant was produced to determine whether the proximity labeling protocol could be used to confirm the role of the N-terminal domain in facilitating homo-oligomerizing interactions to control extracellular binding activities [18, 19]. We chose APEX2 over other peroxidase enzymes, due to its monomeric nature, small size (28 kDa), and improved activities in various cellular compartments [16], especially for proteomic applications [20]. We fused APEX2 to the N-terminus of galectin-3, as previous work has shown that such fusion proteins remain active towards binding glycans [21]. Both fusion proteins were generated efficiently (**Figs. S1A**-**D**) in recombinant *E. coli* and retained their capacity to bind glycans in an Enzyme-Linked Immunosorbent Assay (ELISA) with immobilized asialofetuin (**Fig. S2**). Co-incubation with a soluble competitor, lactose (Lac; Gal-Glc) efficiently blocked binding (**Fig. 2B**), indicating that interactions were glycan-dependent. Apparent binding affinity constants determined for PX-Gal3 (EC_50_ ∼100 nM) versus PX-Gal3_Δ116_ (EC_50_ ∼140 nM) were similar to each other, consistent with prior reports of galectin-3 fusion proteins [21],

**Figure 2.**
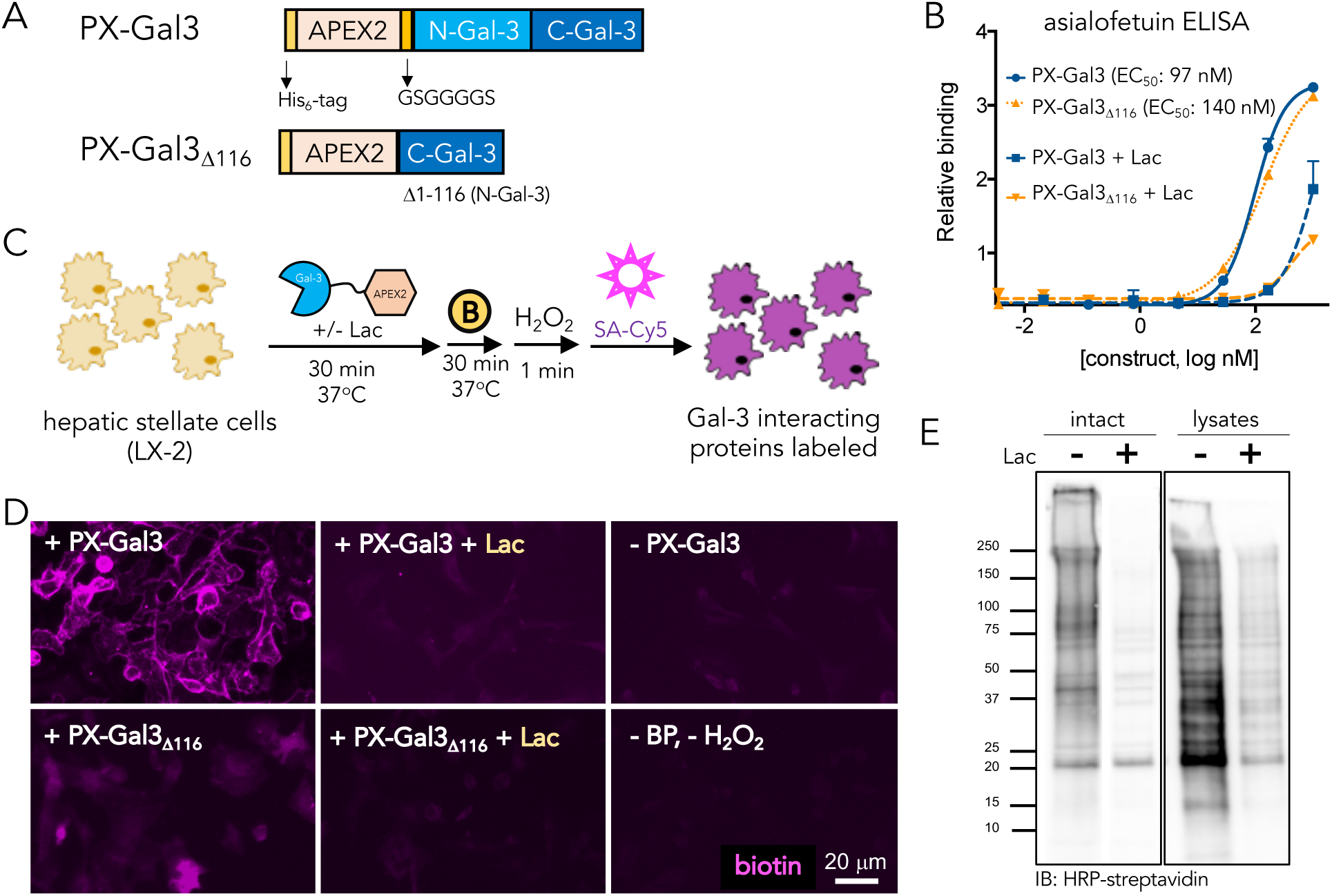
Design and characterization of PX-Gal3 fusion constructs for proximity labeling in LX-2 hepatic stellate cells (HSCs). **(A)** APEX2 fusion constructs of galectin-3, PX-Gal3 and PX-Gal3Δ116, were constructed. The proteins include an N-terminal His-tag sequence, followed by the APEX2 protein, and either full-length galectin-3 or the C-terminal domain of galectin-3. PX-Gal3Δ116 lacks the N-terminal domain of galectin-3, previously determined to be responsible for its homo-oligomerization upon binding glycans at the cell surface. **(B)** Using an ELISA with asialofetuin as a model glycan-bearing ligand, the fusion constructs were determined to retain glycan-binding activities with similar binding affinities (EC50 values). **(C)** Application of the proximal labeling method to live adherent HSCs. The recombinant fusion proteins were first incubated with HSCs (30 min, 37°C). Upon washing to remove unbound proteins, biotin phenol (yellow circle with a B symbol; 1 mM, 30 min, 37°C) was added, followed by hydrogen peroxide (H2O2, 1 min). Biotinylated interactors (purple) were subsequently probed using Cy5-labeled streptavidin (SA-Cy5). **(D)** Fluorescence microscopy images of biotin-tagged (purple) HSCs show that PX-Gal3 (100 nM; 30 min) generates significant labeling over negative controls wherein a component (e.g. protein, or biotin phenol and H2O2) of the labeling protocol is omitted. Co-incubation with lactose (Lac; 100 mM) causes a loss of signal, suggesting that the majority of labeled interactions are glycan-dependent. PX-Gal3Δ116 fails to significantly label interactors. **(E)** Western blotting of biotinylated proteins produced from the proximity labeling method applied in intact live cells versus cell lysates shows that only a subset of interactors is captured.

To examine whether the proximity labeling protocol could identify interactors of galectin-3, we incubated the recombinant fusion proteins with live adherent HSCs. After washing and the step-wise application of biotin-phenol and H_2_O_2_, biotin-tagged interactors were visualized using a fluorophore-conjugated streptavidin probe (**Fig. 2C**) and fluorescence microscopy. We observed significant fluorescence in cells incubated with PX-Gal3 compared with negative controls and PX-Gal3_Δ116_ (**Fig. 2D**), with signals detectable as early as five minutes of protein incubation, which did not significantly increase even up to two hours (**Fig. S3A**). We also observed a dose-dependent increase in fluorescence with increasing amounts (25 to 100 nM) of PX-Gal3 (**Fig. S3B**). In contrast to the plate-based binding assay, PX-Gal3_Δ116_ did not elicit significant signal compared to PX-Gal3. We attribute this observation to the differing glycan density and presentation found in live cells, establishing an important role for the N-terminal oligomerization domain not directly apparent by plate-based methods. This protocol enabled the sensitive detection of interactors with nanomolar concentrations of PX-Gal3, whereas querying for galectin-3 binding to HSCs in simple binding experiments failed to achieve significant signal even at 10 μM concentrations and prolonged incubation periods (**Fig. S3C**). While fluorescence signals were mostly observed at the cell surface, some intracellular labeling was also observed, which increased with protein incubation time (**Fig. S3A**) and PX-Gal3 concentrations (**Fig. S3B**).

Both the co-incubation of Lac (**Fig. S3D**) and a PX-Gal3 mutant that includes three point mutations (R144S, R186A, G182A) that abolish glycan-binding activity [22] failed to show significant fluorescence (**Fig. S4**), indicating that glycan-mediated interactions are highly represented between PX-Gal3 and HSCs. We also compared labeling live intact cells with labeling cellular lysates and found that only a limited subset of potential interactors was captured and substantially blocked with excess Lac (**Fig. 2E; Fig. S5**). Altogether, these observations indicate that the proximity labeling strategy can probe for glycan-dependent interactors in live cells with high sensitivity and that native glycan presentation is important for the selective capture of interactions.

### Identification of biotinylated interactors with quantitative MS-based proteomics

To identify the putative galectin-3 interactors, we subjected HSCs to proximity labeling, followed by cell lysis, streptavidin bead enrichment, and on-bead trypsinization (**Fig. 3A**). We then subjected the resulting peptides to tandem mass tagging (TMT), which chemically tag the N-termini and free amines of the resulting peptides from each condition with a unique isobaric tag that can be distinguished at the MS3 stage [23, 24]. This method allows for the reliable comparative analysis of multiplexed samples in a single run, further ensuring the accuracy of the comparison.

**Figure 3.**
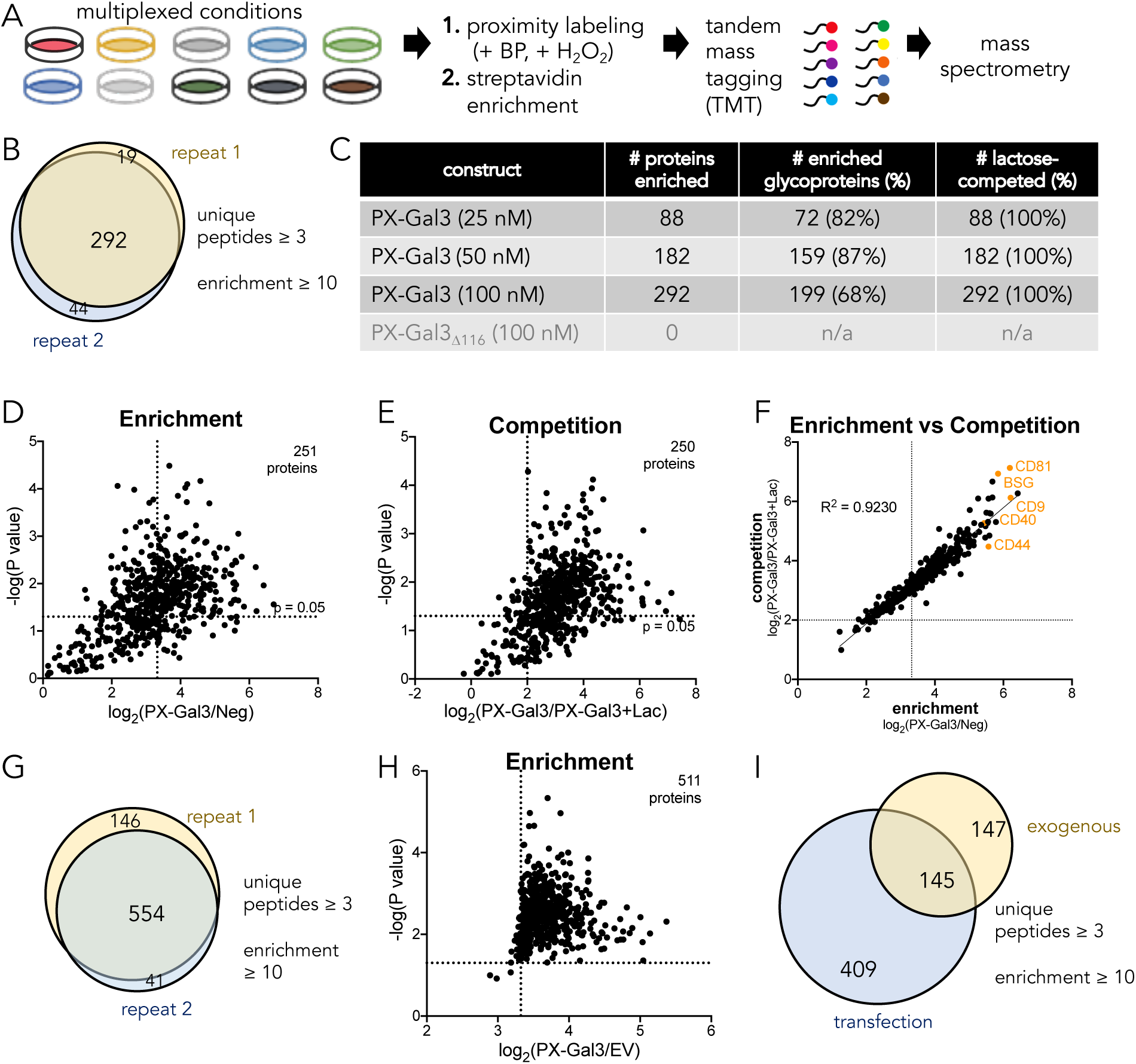
Identification and analysis of the PX-Gal3 interactome by quantitative MS-based proteomics. **(A)** Experimental workflow for the preparation of samples for analysis from LX-2 HSCs. (**B**) 292 proteins were enriched across two biological replicates, defined as proteins with ≥ 3 unique peptides and with TMT ratios (PX-Gal3/Neg) ≥ 10. Neg depicts conditions where the cells were not incubated with any protein, but were still treated with biotin phenol and H2O2. (**C**) Analysis of proteins enriched by PX-Gal3 by dosage, glycosylation status (assigned by UniProt), and competition with lactose. (**D**) Statistically-significant (p < 0.05) and enriched proteins found in cells treated with PX-Gal3 (100 nM, 30 min). (**E**) Proteins that were statistically-significant (p < 0.05) and competed (TMT ratio of PX-Gal3/PX-Gal3+Lac ≥ 4) upon co-incubation with exogenous lactose (100 mM). (**F**) There is high linear correlation between proteins that were enriched and competed by lactose. (**G**) 554 proteins were found to be enriched across two biological replicates when PX-Gal3 is transiently overexpressed in HSCs. (**H**) 511 proteins were found to be significantly enriched (p < 0.05). (**I**) There is some overlap between proteins enriched by the exogenous and transfection protocols.

We identified a total of 292 proteins across two biological replicates (**Fig. 3B; Table 1; Appendix Table S1**). These proteins were consistently detected with ≥ three unique peptides and they were highly enriched (TMT ratio ≥ 10) by PX-Gal3 (100 nM) over the negative control, cells which were not treated with any fusion protein but were similarly treated with biotin phenol and H_2_O_2_ (**Fig. 3B**). Proteins that were non-specifically selected, such as PC, ACACA, MCCC1, were found in the pre-filtered data sets, but these did not fulfill the stringent criteria used to define enrichment (**Fig. S6A**). Notably, we observed a dose-dependent increase in the number of enriched proteins using PX-Gal3 (**Fig. 3C**) as well as an increase in the level of enrichment, whereas no proteins were enriched by PX-Gal3_Δ116_. A majority (68-87%) of enriched proteins in each dose of PX-Gal3 was previously reported to be N-or O-glycosylated (**Fig. 3C**).[25] Among the 292 proteins, 251 (86%) were defined as statistically significant (p < 0.05; **Fig. 3D**); amongst these, 250 (99%) were competed by Lac (TMT Ratio ≥ 4; **Fig. 3E**). There was a high linear correlation (R^2^ = 0.9230) between enriched proteins and Lac-competed interactors (**Fig. 3F; Fig. 3C**). Neither PX-Gal3_Δ116_ (**Fig. S6B**) nor the triple-point mutant protein (**Fig. S6C**) enriched proteins over the negative control. Proteins that were enriched by PX-Gal3 were also competed by TD139 [26], a Lac analogue that displays potent galectin-3 inhibition (**Fig. S6D**).

**Table 1.** Abbreviated list of galectin-3 interactors determined by proximity labeling and quantitative MS spectrometry. An expanded list is available in **Appendix Table S1**. The glycosylation states are assigned according to UniProt [25]. PM: plasma membrane; ERS: Extracellular Region or secreted; Nuc: nucleus; Endo: endosome; Lyso: lysosome

| Gene name | Protein name | UniProt ID | Glycosylation state | Subcellular location |
| --- | --- | --- | --- | --- |
| **BSG** | Basigin | P35613 | 3 N-linked sites | PM; ERS |
| **NPTN** | Neuroplastin | Q9Y639 | 6 N-linked sites | PM |
| **JAM3** | Junctional adhesion molecule C | Q9BX67 | 2 N-linked sites | PM, ERS |
| **SLC35F2** | Solute carrier family 35 member F2 | Q8IXU6 | Not known | PM |
| **SLC7A5** | Large neutral amino acids transporter small subunit 1 | Q01650 | Not known | PM; Lyso |
| **SLC23A2** | Solute carrier family 23 member 2 | Q9UGH3 | 2 N-linked sites | PM |
| **ITGA1** | Integrin-α-1 | P56199 | 26 N-linked sites | PM; ERS |
| **ITGB1** | Integrin-β-1 | P05556 | 12 N-linked sites | PM; Endo |
| **CD44** | CD44 antigen | P16070 | 9 N-linked sites | PM |
| **CD47** | Leukocyte surface antigen CD47 | Q08722 | 6 N-linked sites | PM |
| **CD81** | CD81 antigen | P60033 | Not known | PM |
| **CD9** | CD9 antigen | P21926 | 2 N-linked sites | PM; ERS |
| **TSPAN6** | Tetraspanin-6 | O43657 | 1 N-linked site | PM; ERS |
| **EFNB1** | Ephrin-B1 | P98172/ | 1 N-linked site | PM, Nuc |
| **EFNB2** | Ephrin-B2 | P52799 | 2 N-linked sites | PM |
| **VASN** | Vasorin | Q6EMK4 | 5 N-linked sites | ERS |
| **HNRNPH3** | Heterogeneous nuclear ribonucleoprotein H3 | P31942 | Not known | Nuc |
| **HNRNPD** | Heterogeneous nuclear ribonucleoprotein D0 | Q14103 | Not known | Nuc |
| **HNRNPF** | Heterogeneous nuclear ribonucleoprotein F | P52597 | Not known | Nuc |

Among the PX-Gal3 targets identified by the proximity labeling method (**Appendix Table S1**) are vasorin (VASN), basigin (BSG/CD147), neuroplastin (NPTN), members of the tetraspanin family (CD9, CD81, TSPAN6), members of the ephrin family (EPHB1, EPHB2), members of the solute carrier family of transporters (SLC35F2, SLC23A2, SLC7A5), and members of the integrin family (ITGA1, ITGB1). Although many of the targets are assigned to plasma membrane or cell surface locations, several are also found intracellularly, including members of the human ribonucleoprotein (HNRNP) complex [27], which are found in the nucleus. These results indicate that despite their exogenous application, the fusion proteins traverse pathways that are expected of the dynamic nature of galectin-3 - first interacting with cell surface proteins and then eventually traversing the cell membrane to associate with intracellular components [28].

We derived effective galectin-3 binding affinity constants (EC_50_) for select proteins (basigin, CD47, ITGA1, JAM3, ephrin-B2, vasorin) using the proximity labeling protocol and TMT quantitation (**Fig. S6E**). Such binding constants are reflective of native glycan and protein expression levels. While it is difficult to extrapolate these constants as a comparative measure of binding of galectin-3 against individual proteins, it is gratifying to see that the proximity labeling method can achieve robust nanomolar binding against these proteins in cells. We used Integrated Pathway Analysis (IPA) to derive putative binding relationships and observed that basigin lies at a nexus towards other proteins that were enriched by PX-Gal3, such as integrin-β-1, SLC7A5, JAM3, and SLC23A2 (**Fig. S6F**).

Considering that we identified intracellular protein targets with exogenously administered PX-Gal3, we set out to evaluate whether proximity labeling could also identify interactors for galectin-3 when it is expressed intracellularly. Thus, we overexpressed PX-Gal3 in HSCs by transient transfection. (**Fig. S7A, B**). Following transfection, some cells were further treated with tunicamycin, an N-glycosylation inhibitor, to assess whether N-linked glycosylation played a role (Lac is impermeable to cells). A total of 554 proteins (filtered by ≥ 3 unique peptides) were found to be enriched (TMT ratio over empty vector control ≥10) and overlapping across two biological replicates (**Fig. 3G; Appendix Table S1**), 511 (92%) of which were statistically significant (p < 0.05; **Fig. 3H**). Interestingly, treatment with tunicamycin (**Fig. S7C**) failed to compete (TMT ratio ≥ 4) for labeling, suggesting that intracellular galectin-3 interactions are mostly mediated by protein-protein rather than protein-glycan interactions, consistent with previous reports [28]. Comparing the enriched proteins from the exogenous treatment (**Fig. 3B**) versus the transient over-expression of PX-Gal3 yielded 145 proteins in common (**Fig. 3I**), possibly suggesting that galectin-3 may associate with some proteins both in a glycan-mediated or non-glycan-mediated manner, depending on its entry.

### Validation of interactions between galectin-3 and identified proteins

As the proximity labeling strategy only tags proteins (< 20 nm) that occur within the vicinity of galectin-3 and these may not necessarily represent direct receptors, we next sought to examine whether selected proteins identified by the proteomic experiments are direct receptors for galectin-3. We confirmed that basigin, CD9, ephrin-B1, CD47, and vasorin are prominently expressed in LX-2 cells (**Fig. S8**). Using immobilized human recombinant proteins (**Table S1**) of basigin, CD9, CD47, CD81, ephrin-B1, neuroplastin, and vasorin, we found that galectin-3 binds in a dose-dependent manner (**Fig. 4A; Table S2**). Apparent binding affinities, with EC_50_ values ranging from 0.7 μM (CD9) to 4.2 μM (CD47) were observed (**Table S2**). Notably, we found that the majority of the proteins were competed by the presence of either Lac or TD139, CD81 binding was not blocked by either competitor, and CD9 interactions could only be competed by TD139 (**Fig. 4B**). Human CD81 is not known to be glycosylated, suggesting that its binding to galectin-3 might be mediated by secondary protein-protein interactions. Human CD9 (UniProt P21926) is predicted to have two N-linked glycosylation sites, implicating potential glycan-mediated interactions with galectin-3, but that these interactions may contribute only partially to galectin binding to CD9, as competition with TD139 was still observed. Altogether, these results indicate that basigin, CD9, CD47, CD81, ephrin-B1, vasorin, and neuroplastin, are potential receptors for galectin-3 in cells.

**Figure 4.**
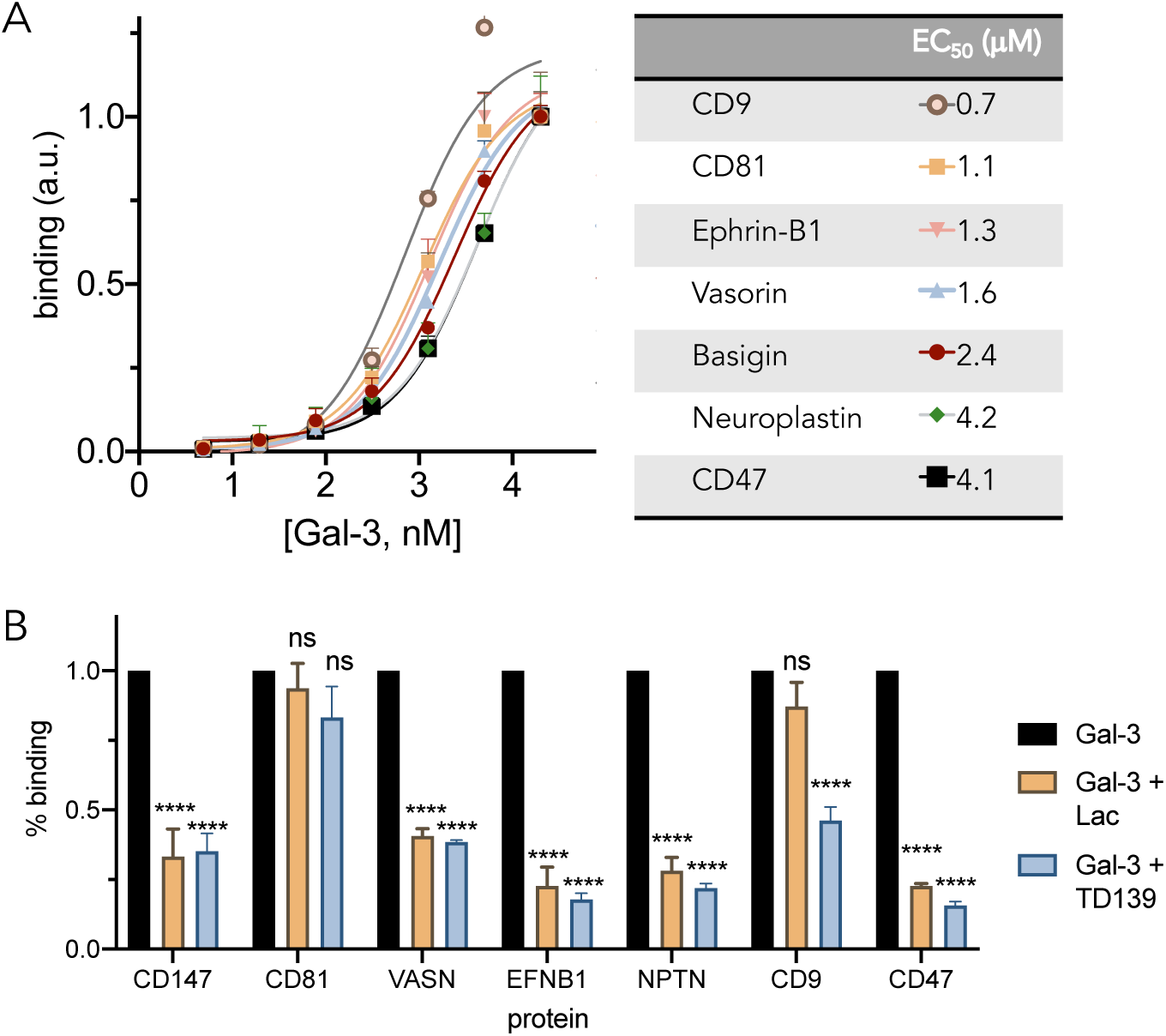
Validation of binding between galectin-3 and proteins identified from proteomics analysis. (**A**) ELISA binding curves and corresponding EC50 values determined for recombinant proteins against galectin-3. **(B)** Co-incubation of galectin-3 (5 μM) with lactose (Lac, 100 mM) or TD139 (15.4 μM) competes for glycan-mediated binding interactions in ELISA.

### MS-based characterization of glycomes from LX-2 cell surfaces, PX-Gal3 enriched interactors, and endogenous basigin

Given the importance of cell surface glycosylation in mediating galectin-3 interactions (**Figs. 2 and 3**), we next sought to identify the structures of the glycans found in LX-2 cells. While several groups have reported that O-linked glycans from mucins [29] and glycosaminoglycans [30] can serve as ligands for galectin-3, complex galactose-terminating N-glycans (with some tolerance to sialylation) have been definitively established as the prominent high-affinity binding galectin-3 epitopes in cells [31, 32]. Thus, we next focused our investigations towards profiling N-glycans, with particular attention towards assessing the abundance of N-glycans bearing the high-affinity terminal galactose (yellow circle) monosaccharides [33]. We further evaluated whether certain glycans were enriched by PX-Gal3 and whether they were present on one of the identified glycoprotein receptors for galectin-3 (**Figs. 3 and 4**).

We harvested N-glycans from intact LX-2 cells by brief treatment with trypsin and PNGase F. The free N-glycans were then subjected to reduction and permethylation, prior to MS analysis. Using the extracted ion intensities acquired during the full MS scan, relative quantitation of the most abundant N-glycans was determined [34]. The most abundant N-glycans found on LX-2 cell surfaces consist mostly of complex and oligomannose N-glycans, with the most abundant being the di-sialylated FA2G2S2 N-glycan, comprising ∼20% of the total population (**Fig. 5; Fig. S9; Fig. S10**). LX-2 cells stained positively with *Sambucus nigra* and *Maackia amurensis* lectins, indicating that both α(2-3) and α(2-6) terminal Neu5Ac sialic acids, respectively, are present (**Fig. S11**). Oligomannose N-glycans (M7, M6, M8) were the next most abundant N-glycans, representing a cumulative total of ∼27% of the cell surface N-glycome. Terminal mono- and di-galactosylated N-glycans, FA2G1 and FA2G2, only comprised ∼3% and ∼1% of the population, respectively.

**Figure 5.**
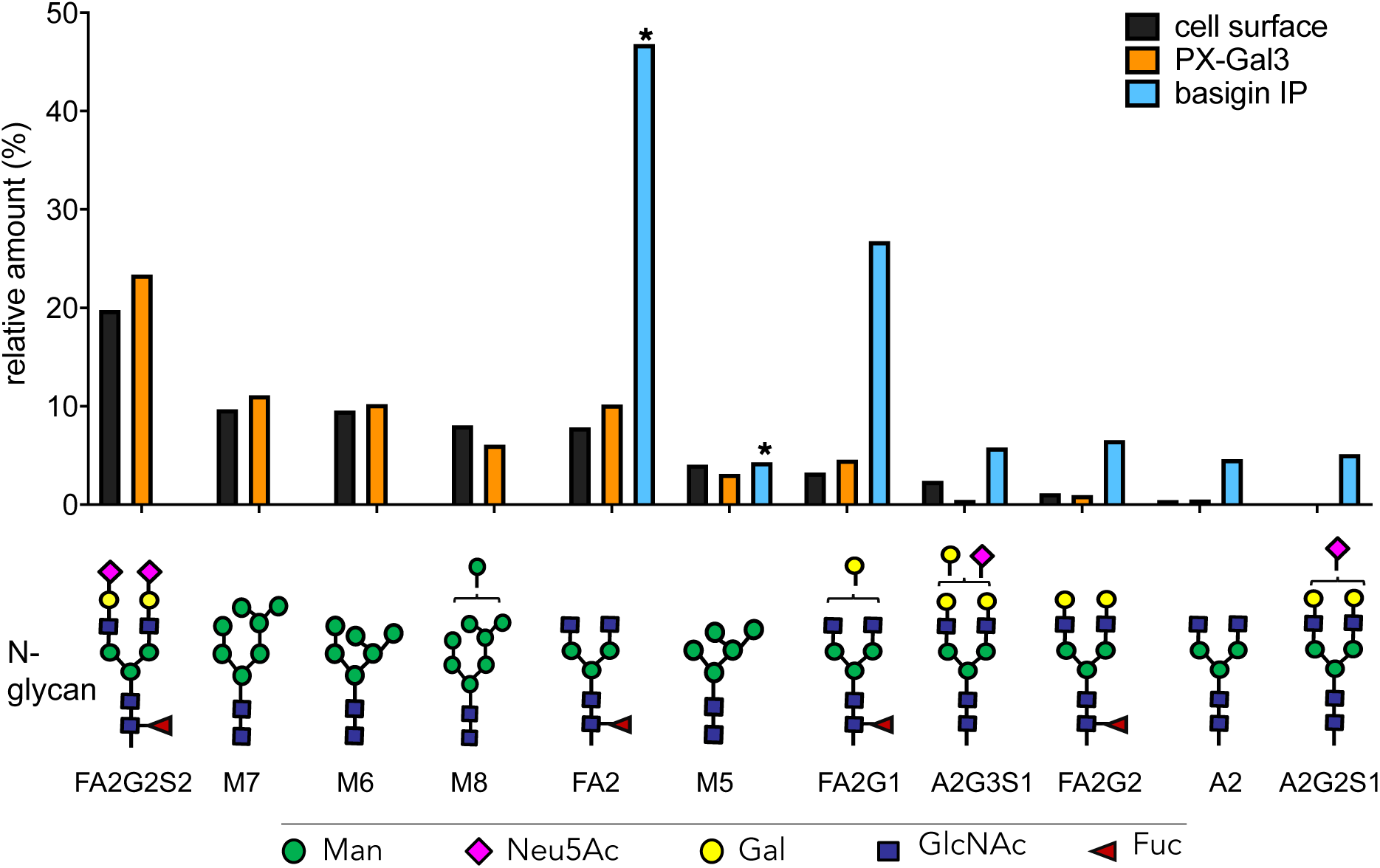
MS-based determination of N-glycan abundance and composition. Composition and relative amounts of the most abundant N-glycans found on LX-2 cell surfaces (black bars), glycoproteins enriched by PX-Gal3 (orange bars), and immunoprecipitated (IP) basigin (blue bars). The most abundant N-glycans found in basigin sample greatly differ from the cell surface samples. The asterisk (*) symbol indicates N-glycans which were also found in the antibody used for IP. Terminal galactose (yellow circles) presenting N-glycans were found to be augmented in basigin compared to the cell surface. Oligomannose glycans bear terminal mannose residues (green circles) and are designated with an M (M7, M6, M8, M5), all others shown here are complex type N-glycans.

To identify N-glycans from the resulting mixture of peptides and glycopeptides captured from proximity labeling with PX-Gal3 (**Fig. 2A**), these samples were also subjected these to PNGAse F treatment. The most abundant N-glycans from these samples were overall quite similar to the ones found on LX-2 cell surfaces (**Fig. 5; Fig. S12**). FA2G1 and FA2G2 composed ∼5% and ∼1% of the total N-glycome, respectively. We observed a modest growth in the amount of terminal galactosylated N-glycans, with the di-galactosylated FA3G2 showing a ∼57% increase compared to the cell surface (**Fig. S13**).

Given the central role of basigin as a glycan-dependent binding receptor for galectin-3 (**Fig. 4**) and its ability to bind other enriched proteins (**Fig. S6F**), we sought to determine its native N-glycosylation pattern. Towards this end, we immunoprecipitated basigin from LX-2 cells using an anti-basigin antibody and protein A beads (**Fig. S14**) and observed pronounced differences compared to the cell surface N-glycome (**Fig. 5**). In contrast to the cell surface, which was composed of a mixture of complex and oligomannose N-glycans, the six most abundant N-glycans were found to be complex. The complex N-glycan FA2 composed ∼47% of the total population, although this N-glycan was also found in the anti-basigin antibody (**Fig. S15**). The next most abundant glycans exclusively found in this sample were found to be the mono- and di-galactosylated FA2G1 and FA2G2, comprising ∼27% and ∼7%, respectively. The mono-sialylated A2G3S1 and A2G2S1 were also abundant in this sample. These observations confirm that the high-affinity galactose epitope is present on LX-2 cell surfaces and that it is highly abundant on basigin.

## Discussion

Our overall goal is to determine whether a proximity labeling approach would be suitable to map the glycan-mediated interactors for galectin-3 in live cells (**Fig. 1**). Here, we have demonstrated that a one minute live labeling step with a fusion protein of APEX2 and galectin-3 covalently tagged protein interactors in hepatic stellate cells (**Fig. 2**). Integration of this method with quantitative MS-based proteomics identified 292 enriched proteins, majority of which are glycosylated (**Fig. 3**). Prior to this report, only 100-185 proteins were found in complex with galectin-3 using standard affinity techniques [5-10], and our dataset extends the number of previously known interactors for galectin-3. These enriched proteins relied exclusively on glycan-dependent interactions for catalytic tagging. In contrast to in vitro experiments and tagging procedures on lysed cells, we showed that the N-terminal oligomerization domain [19] is critical for productive cell surface interactions (**Fig. 2D**), and despite many other possible interactors (**Fig. 5**), only a subset form favorable interactions with galectin-3 (**Fig. 2E**). While other labeling strategies use horse radish peroxidase fusion proteins [35] that require oxidative conditions, our method uses the APEX2 enzyme for covalent tagging. APEX2 is active in both oxidative extracellular conditions and the reducing environment of the intracellular milieu [36], permitting the identification of 554 interactors for PX-Gal3 when it is intracellularly over-expressed (**Fig. 3**). Consistent with previous reports, we also found that intracellular interactions with galectin-3 are not glycan-dependent [28].

Informed by MS-based proteomics, we selected seven proteins to confirm direct interactions with galectin-3, and found that recombinant basigin, CD9, CD47, CD81, ephrin-B1, neuroplastin, and vasorin bound galectin-3, and binding to CD9 and CD81 were not glycan-mediated in vitro (**Fig. 4**). It must be noted however, that these recombinant proteins may not necessarily reflect the native protein sequence or glycosylation patterns found in LX-2 cells (**Table S1**). CD47, ephrin-B1, and vasorin are previously unreported glycan-mediated ligands for galectin-3. While a majority of proteins enriched by the proximity labeling strategy are glycoproteins (**Fig. 3C**), some are not known to possess sites for N-linked glycosylation. These observations highlight that while glycan-dependent interactions are responsible for initial binding events with galectin-3, other protein-protein interactions also occur subsequent to the primary interaction. Thus, additional experiments are required to definitively assign the direct glycoprotein ligands for galectin-3 within the 292 proteins.

Both CD9 and CD81 are members of the tetraspanin family, a group of proteins that organize other proteins into discrete membrane compartments to regulate trafficking and signaling [37]. CD9 and CD81 have been found in complex with each other and with other glycoproteins [38, 39], and it is intriguing to surmise how membrane organization could play a role in facilitating downstream signaling events upon the binding of galectin-3 to HSCs. We speculate that galectin-3 binding either induces clustering of glycoproteins or it may bind pre-formed clusters in HSCs, and that such organized microdomains could similarly promote galectin-3-dependent signal transduction events [29] that lead to HSC activation [40].

Changes in LX-2 cell surface glycosylation [40] and upregulation of terminal galactosylation are associated with HSC activation [41]. Using MS-based glycomics, we determined that LX-2 cell surfaces are mostly composed of complex and oligomannose N-glycans (**Fig. 5**). Notably, the N-glycome of endogenous basigin substantially differs from that of the cell surface, and the terminal galactose epitope is highly represented. These observations validate why LX-2 cell surfaces and basigin can serve as ligands for galectin-3. Although basigin has not been shown to display terminal galactosides [42, 43], such differences may reflect the natural abundance of glycan biosynthetic enzymes and sugar nucleotide precursors between different tissues. Human basigin naturally occurs in four natural isoforms [44], and our proteomics data suggests that either basigin-1 or basigin-2 is expressed in LX-2s. Endogenous basigin-2 differs from the canonical basigin-1 by missing regions 24-139 (**Fig. S16**) and similarly bears three N-linked glycosylation sites. The recombinant basigin glycoprotein used for the *in vitro* binding experiments only possessed two sites for N-glycosylation (**Fig. S16**), suggesting that tighter interactions may even be achieved by the endogenous glycoprotein.

Empowered by the proximity labeling approach, we now have generated a priority list of proteins that will be the subject of future work to further reveal the “professional glycoprotein ligands” [14] required for the functional activation of HSCs. Already, preliminary work has shown that basigin plays an active role in hepatic fibrosis [45], and our own work will delve deeper into the molecular nature of the galectin-3-basigin interaction. The glycoproteomic assignment of which N-glycans are presented on individual Asn sites, and the identification of which of the three N-glycosites can bind galectin-3, as well as connecting the galectin-3-basigin interaction nexus and its functional consequences in signaling and HSC differentiation, are of particular importance. The PX-Gal3 fusion constructs and the proximity labeling approach could easily be extended towards studies in other biological samples, such as primary cells and tissues, and they could also be used to track the spatiotemporal dynamics of galectin-3 interactions [46] or to visualize interactions using electron microscopy [17].

## Conclusions

GBP-glycan interactions choreograph many biological events, yet they are often interrogated in non-native environments without the identification of the glycan-bearing glycoprotein receptors. Herein, we used a proximity labeling method consisting of fusion proteins of galectin-3 and the APEX2 enzyme to map glycan-mediated galectin-3 interactions in live cells. This method is sensitive, enabling detection of galectin-3 binding at nanomolar concentrations, and we have produced an inventory of proteins that occur in complex with galectin-3, and some of these proteins were validated as direct glycan-mediated glycoprotein receptors. Consequently, our results provide greater, higher-resolution insight into the glycan and protein determinants that govern galectin-3 interactions in hepatic stellate cells and with individual glycoproteins. Critically, although we have mapped the galectin-3 interactome with hepatic stellate cells, the functional consequences that these discrete interactions have on hepatic fibrosis remain unknown. Informed by our results, we are now poised to formulate new hypotheses that define the contributions of individual proteins and glycosylation sites. Importantly, we expect that the proximity labeling approach is generally applicable towards other GBP and glycoproteins, and that it will be a powerful tool to advance our understanding of glycan-mediated cell biology.

## Experimental Procedures

### Generation of PX-Gal3 fusion constructs

The full-length DNA insert for PX-Gal3 and PX-Gal3_Δ116_ were synthesized by Genscript (New Jersey, USA). Starting from the human galectin-3 cDNA sequence, an optimized sequence that has a codon adaptation index of 0.96 was determined by Genscript using a proprietary OptimumGene algorithm and ligated into the EcoRI/NotI cloning sites of a pETDuet-1 vector. Upon transformation into BL21 (DE3) pLysS chemically competent *E. coli* and antibiotic selection using ampicillin LB agar plates (100 μg/mL), single cell colonies were picked, and the selected clones grown in LB with ampicillin (100 μg/mL). To induce protein expression, the transformed E. coli was grown in 5 mL starter cultures of LB with ampicillin overnight. The overnight culture was then added to 500 mL of fresh LB with ampicillin and shaken at 37°C until OD_600_∼0.6. The cultures were then induced with IPTG (1 mM, 4 hours at 37°C) to initiate protein expression in LB containing ampicillin (100 μg/mL). Following centrifugation (3300 xg, 15 min at 4°C) and lysis of the cell pellet in lysis buffer on ice (5 mL; 20 mM imidazole in 2x PBS (274 mM NaCl, 5.4 mM KCl, 20 mM Na_2_HPO_4_, 3.6 mM KH_2_PO_4_) *via* sonication (1 sec ON, 3 sec OFF, 30 sec TOTAL ON, 20% amplitude), the crude protein supernatant was purified using Ni-NTA beads (1 hr at 4°C) or an AKTA Start FPLC system for His-tag mediated purification. The column-bound proteins were washed with 25 mM imidazole and eluted using 250 mM imidazole, and further purified using α-lactose agarose beads. Upon elution with 250 mM lactose, the resulting proteins were dialyzed against PBS to remove excess lactose and stored in PBS supplemented with 2mM EDTA and 0.05% Tween-20, flash frozen and stored at-80°C. SDS-PAGE and immunoblotting were used to verify expression and purity.

### Growth and maintenance of hepatic stellate cells

LX-2 human hepatic stellate cells [47] (Millipore) were cultured in DMEM supplemented with 10% fetal bovine serum at 37°C under 5% CO_2_. For fluorescence microscopy imaging experiments, cells were grown on plastic 24-well culture plates. To improve the adherence of LX-2 cells, dishes were pre-coated with poly-D-lysine and washed with DPBS prior to cell plating.

### In situ proximity tagging in live LX-2 cells and imaging

The LX-2 cells were plated (pre-coated with poly-D-lysine) at a density of 1.5 x 10^5^ cells per well of a 24-well plate plastic dish overnight at 37°C, 5% CO_2_. The next day, the cells were gently washed with DPBS and incubated with DMEM supplemented with the fusion protein of interest for defined periods of time (usually 30 min, unless otherwise indicated) at 37°C, 5% CO_2._ After washing twice with DPBS to remove unbound proteins, labeling was initiated by replacing the media with pre-warmed 10% FBS/DMEM containing 500 μM biotin-phenol. This solution was further incubated with the cells at 37°C, 5% CO_2_ for 30 min. H_2_O_2_ was then added to each well to achieve a final concentration of 1 mM, and the plate was gently agitated for 1 minute at room temperature (RT). After aspirating the solution, the reaction was then quenched by washing three times with quencher solution (5 mM Trolox, 10 mM sodium ascorbate, and 10 mM sodium azide in DPBS). Cells used for fluorescence imaging were then fixed with 4% p-formaldehyde in PBS (RT, 10 min) and washed with PBS. To probe for biotinylated interactors, Cy5-streptavidin was used in PBST to incubate (1 hr, dark, RT). Imaging was performed on a benchtop EVOS M5000 (ThermoFisher, Carlsbad, CA) fluorescence imager. Experiments were repeated with at least two independent biological replicates.

### Preparation and enrichment of sample lysates for proteomics

LX-2 cells were plated onto a single dish per condition. Cells were plated (pre-coated with poly-D-lysine) at a density of 8-10 x 10^6^ cells per plate overnight at 37°C, 5% CO2. The next day, the cells were gently washed with DPBS and incubated with DMEM supplemented with the fusion protein of interest for defined periods of time (usually 30 min, unless otherwise indicated) at 37°C, 5% CO2. After washing twice with DPBS to remove unbound proteins, labeling was initiated by replacing the media with pre-warmed 10% FBS/DMEM containing biotin-phenol (500 μM). This solution was further incubated with the cells (37°C, 5% CO2, 30 min). H2O2 was then added to each well to achieve a final concentration of 1 mM, and the plate was gently agitated for 1 minute at room temperature (RT). After aspirating the solution, the reaction was then quenched by washing three times with quencher solution (5 mM Trolox, 10 mM sodium ascorbate, and 10 mM sodium azide in DPBS). Cells were then washed with DPBS, scraped and transferred to 15 mL falcon tubes, pelleted and stored at -80°C until the next step. Cell pellets were resuspended in 300-400 μl DPBS and lysed by sonication (15 ms ON, 40 ms OFF, 1 s total ON, 15% amplitude x2). Protein concentrations were normalized (1.5 – 2 mg/mL; final volume of 500 μL with PBS) using the 660nm Protein Assay (Pierce). Each lysate was transferred to a new 15mL falcon tube and cold mixture of MeOH:CHCl3 (4:1) (2.5 mL) was added to the lysate followed by cold DPBS (1 mL). The resulting mixture was vortexed followed by centrifugation (5,000 RPM, 10 min, 4°C). The organic and aqueous layers were aspirated and the remaining protein disc was further washed via sonication in cold MeOH:CHCl3 (4:1) (2mL) and pelleted by centrifugation (5,000 RPM, 10 min, 4°C). The protein pellet was aspirated and dissolved in 500 μL of freshly-prepared proteomics-grade urea (6 M in DPBS) with 10 μL of 10% SDS by sonication. Disulfide bonds were reduced by adding 50 μL of freshly prepared 1:1 solution of TCEP (200 mM in DPBS) and K2CO3 (600 mM in DPBS) for 30 mins at 37°C on a shaker. Free thiols were alkylated by adding 70 μL of freshly prepared iodoacetamide (400 mM in DPBS) at RT in the dark. To each solution, 130 μL of 10% SDS in DPBS was added and each sample was diluted with DPBS (5.5 mL) and incubated with pre-equilibrated streptavidin-agarose beads (100 μL of 50% slurry, Pierce) for 1.5 hr at RT while rotating. The streptavidin beads were pelleted by centrifugation (2,000 RPM, 2 min, 4°C) and sequentially washed with 0.2% SDS in DPBS (1 x 5 mL), DPBS (2 x 5mL), and 200 mM EPPS pH 8.4 (1 x 5mL). The beads were transferred into LoBind Microcentrifuge Tubes (Eppendorf # 022431081) and bound proteins were digested for ∼14 hr at 37°C in 200 μL of trypsin premix solution containing sequence grade trypsin (2 mg, Promega), urea (0.5 M in 200 mM EPPS pH 8.4), and calcium chloride (1 mM in MQ H2O). The beads were pelleted by centrifugation (2,000 rpm, 2 min, 4°C) and the supernatant containing the digested peptides were transferred to new LoBind Microcentrifuge Tubes (Eppendorf). Each digested sample was labeled with tandem mass tag (Thermo Scientific cat# A34808). In general, for each sample, 8 μL of the 20 μg/μL stock of TMT reagent was added along with dry MS-grade acetonitrile to achieve a final acetonitrile concentration of approximately 30% (v/v), following incubation at room temperature for 1 hr. The reaction was quenched by adding 6 μL of hydroxylamine for 15 min, followed by acidification with the addition of 5μL of formic acid. Each TMT-labeled sample was dried through vacuum centrifugation and the samples were combined by re-dissolving one sample with Buffer A (400 mL; 5% MeCN in H2O, 0.1% FA) and transferring solution into each sample tube until all samples were re-dissolved (final volume ∼ 600 mL). pH was adjusted to ∼3 with FA. Combined sample was desalted using two C-18 columns (Thermo Fisher # 89870) according to the instruction manual. Sample was dried through vacuum centrifugation and stored in the -80°C until ready for injection.

### LC-MS analysis

10-plex samples were re-dissolved in 20 μL Buffer A (99.9% H2O, 0.1% formic acid) for MS analysis. Briefly, 3 μL of each sample was loaded onto a Thermo Acclaim PepMap100 precolumn (75 μM × 2 mm) and eluted on a Thermo Acclaim PepMap RSLC analytical column (75 μM × 15 cm) using a Thermo UltiMate 3000 RSLCnano system. Buffer A (0.1% formic acid in H2O) and Buffer B (0.1% formic acid in acetonitrile) were used to establish the 220 min gradient comprised of 10 min of 2% B, 192 min of 2-30% B, and 5 min of 30-60% B, 1 min of 60-95% B, 5 min of 95% B, 1 min of 95-2% B, followed by re-equilibrating at 2% B for 6 min. The flow rate was 0.3

μL/min at 35°C. Peptides were than analyzed on a Thermo Orbitrap Fusion Lumos proteomic mass spectrometer in a data-dependent manner, with automatic switching between MS and MS/MS scans using a cycle time 3 s. MS spectra were acquired at a resolution of 120,000 with an AGC target value of 1×106 ions or a maximum integration time of 50 ms. The scan range was limited from 375 to 1500 m/z. Peptide fragmentation for MS2 was performed via collision-induced dissociation (CID) with the energy set at 30% collision energy and 10 ms activation time. The detector was Ion trap with an AGC target value of 1.0×104 ions or a maximum integration time of 120 ms. The fixed first m/z was 120, and the isolation window was 1.6 m/z. MS3 precursor was fragmented via high energy collision-induced dissociation (HCD) with the collision energy of 65%. The detector type was orbitrap with a resolution of 50000, an AGC target value of 1.0×105, and a maximum integration time of 105 ms.

### Proteomics data analysis

Data processing was performed using Proteome Discoverer 2.4 software (Thermo Scientific) and peptide sequences were determined by matching protein databases with the acquired fragmentation pattern by SEQUEST HT algorithm. The precursor mass tolerance was set to 10 ppm and fragment ion mass tolerance to 0.6 Da. One missed cleavage site of trypsin was allowed. Carbamidomethyl (C) and TMT-6plex (K and N-terminal) were used as a static modification. Oxidation (M) was used as variable modification. All spectra were searched against the proteome database of homo sapiens (42358 sequences) using a target false discovery rate (FDR) of 1%. The proteins identified were additionally filtered by at least three unique peptides. Statistical analysis was performed with Prism 8 (GraphPad Software Inc., CA). TMT ratios obtained from Proteome Discoverer were transformed with log2-(x), p-values were obtained from t-test function over two biological replicates. Quantitative data are listed in Appendix Table S1. The raw data and analysis files for proteomic analyses were deposited to the ProteomeXchange Consortium (http://proteomecentral.proteomexchange.org) via xxx.

### Gels and Western blots

Unless otherwise stated, SDS-PAGE analysis of proteins was performed using either manually cast or pre-cast protein gels (Biorad). Samples were loaded using Laemmli buffer with 2-mercaptoethanol and boiled (95°C, 5 min). Running conditions include 85-100 mV in Tris-Glycine-SDS buffer for 60-90min. Transfers were performed using nitrocellulose or PVDF membranes (60 min, 100 mA). Blocking was performed with 5% BSA/TBST, incubations with primary antibodies were performed overnight at 4°C, with rocking. Secondary antibody incubations were performed at RT for 1-2 hours.

### ELISAs to evaluate binding affinities

The glycoprotein(s) of interest were immobilized at defined concentrations (specified in each context) in pH 9.6 carbonate buffer overnight at 4°C onto high-binding plate (Nunc Maxisorp). Next day, plates were washed 3x in PBST, followed by blocking with 2% BSA/PBST (1 hr, RT). Recombinant galectin-3 (Biolegend # 599706) was then incubated either for 1-2 hrs at RT or overnight at 4°C, with rocking. Following washing 3x with PBST, plates were incubated with anti-galectin-3 antibody, followed by an HRP-conjugated secondary antibody. Following subsequent washing steps, plates were developed with TMB substrate and quenched with 2N H_2_SO_4_. For ELISAs with asialofetuin (Sigma Aldrich # A4781), 5 μg/mL was immobilized. For ELISAs with multiple targets, equimolar (40 nM) amounts of the recombinant human proteins were immobilized onto the plate. Experiments were repeated with at least two independent biological replicates.

### Transient over-expression of PX-Gal3

Adherent LX-2 cells seeded overnight at 50-80% confluency (1.25 x 10^5^ cells/well) were washed twice in DPBS and incubated with plasmid DNA pre-complexed (according to manufacturer’s instructions) with Fugene HD and returned to the incubator. The DNA sequence encoding PX-Gal3 was inserted into a bicistronic pIRES2-AcGFP1 vector (Takarabio # 632435). GFP fluorescence was readily observed by microscopy 24 and 48 hours after incubation, and subsequent proximity ligation steps were performed as before, while omitting the initial protein incubation step.

### N-glycan glycomics MS of LX-2 cell surfaces

Cell surface N-glycomics analysis was based on a published procedure [48], starting from LX-2 cells. The cells were cultured to ∼ 90% confluency in 10 cm^2^ plates to yield approximately 10 x 10^6^ cells per plate. Cells were harvested and washed four times by re-suspending in 4 mL ice-cold PBS followed by centrifugation (600 rpm, 10 min, 4°C). Free glycopeptides were obtained by re-suspending cell pellets in 1.0 mL of 0.25% trypsin in PBS (Quality Biological, #118086721) and shaking at 250 rpm for 15 min at 37°C. Samples were placed in ice for five min followed by centrifugation (8,000 x*g*, 10 min, 4°C). The resultant glycopeptide-containing supernatant was heated at 100°C for five min to deactivate trypsin. After cooling for 10 min on ice, glycans were released by treatment with 2.0 µL of PNGase F for 20 hr at 37°C (in-house, Addgene #114274; 7.5 µg/µL in 25 mM Tris pH 7.2 with 50% glycerol). Following the treatment with PNGase F, peptides were removed with a C18 cartridge (Thermo Scientific HyperSep, #03251257). Column cartridges were conditioned by successive treatment with methanol, 5% aqueous acetic acid, *n*-propanol, and 5% aqueous acetic acid. Samples were first acidified to 5% acetic acid, centrifuged to remove debris (16,000 x *g*, 10 min, 4°C), then loaded to the column and eluted with 5.0 mL 5% aqueous acetic acid. Column flow-through was combined with aqueous eluate and lyophilized. Experiments were repeated across two independent biological replicates.

### Preparation of PX-Gal3 enriched glycans

Approximately 400 µL of the experimentally-treated beads were suspended in 1.5 mL 0.25% trypsin in PBS (Quality Biological, #118086721) and shaken at 250 rpm for 30 min at 37°C. This was then heated at 100°C for five min to denature the trypsin. After cooling on ice for five min the sample was treated with 5.0 µL of PNGase F for 20 hr at 37 °C (in-house, Addgene #114274; 7.5 µg/µL in 25 mM Tris pH 7.2 with 50% glycerol) and incubated at 37°C for 20 hr. The glycan-containing supernatant was separated from the beads, and peptides were removed with a C18 cartridge (Thermo Scientific HyperSep, #03251257) conditioned by successive treatment with methanol, 5% aqueous acetic acid, *n*-propanol, and 5% aqueous acetic acid. The sample was centrifuged to remove debris (14000 x*g*, 10 min, 4°C), loaded onto the column and eluted with 5.0 mL 5% aqueous acetic acid. The aqueous flow through and eluate were combined and lyophilized.

### Immunoprecipitation of endogenous basigin from LX-2 cells

Confluent plates of 15 cm^2^ plates of live LX-2 hepatic stellate cells were scraped, and cells were pelleted by centrifugation (650 xg, 3 min, 4°C). The cells were washed with cold PBS twice and then lysed with RIPA + Protease Inhibitor Cocktail (1 mL) for 15 min on ice. The lysate was pre-clarified by centrifugation (16,000 xg, 10 min, 4°C) and the supernatant was added to pre-washed Protein G Sepharose Fast Flow beads (100 μL, GE) for 10 min at 4°C on a rocker to remove any non-specific bead binding. The beads were pelleted by centrifugation (1,450 xg, 10 min, 4°C). The immunocomplex was captured by adding the lysate and 10 μg of anti-basigin antibody (Novus Biologicals # NB500-430) concurrently with pre-washed Protein G Sepharose Fast Flow beads (100 μL, GE) overnight at 4°C on a rocker. The beads were pelleted by centrifugation (1,450 xg, 10 min, 4°C) and the supernatant was collected and put aside. The beads were then washed with DPBS (1 mL x 3). Basigin was eluted from the antibody on the beads by adding 100 μL of elution buffer (0.2 M glycine, pH 2.6) for 10 min at RT. The beads were pelleted and the supernatant was removed and quickly quenched with an equal volume of Tris buffer (1 M, pH 9.0). The elution and quenching was repeated two more times, for a total of three elutions and kept separate. 4x Laemmli buffer was added to the beads and heated at 95°C for 10 min to dissociate any remaining immunocomplex off the beads.

### Harvesting N-glycans from immunoprecipitated basigin

To prepare and purify the N-glycans captured by basigin immunoprecipitation, the eluate from the BSG-antibody beads (as well as positive and negative controls) was lyophilized, re-dissolved in 300 uL water, and proteins were denatured by heating at 80°C for 30 minutes. After cooling on ice, the samples were digested with 2.0 µL of PNGase F for 20 h at 37°C (in-house, Addgene #114274; 7.5 µg/µL in 25 mM tris pH 7.2 with 50% glycerol). Free glycans were separated from protein with a 10 kDa molecular weight cutoff centrifugal filter unit (Millipore Amicon Ultra-0.5 #UFC5010). The glycan-containing filtrate was desalted with a PD minitrap G-10 column (Cytiva) and dried under vacuum.

### Preparation of released N-glycans for mass spectrometry analysis

The released N-glycans from the cell surface, PX-Gal3, and basigin-IP samples were prepared for LC/MS analysis by reduction followed by permethylation. Reduction was carried out by dissolving the dried glycans in 200 µL of 10 mg/mL sodium borohydride (Oakwood Chemical # 042896) in 1 M NH_4_OH solution (Sigma-Aldrich # 09859) and heating at 60°C for 1 hr. After cooling to RT, the samples were desalted using a PD minitrap G-10 column (Cytiva) and lyophilized. Permethylation was based on a published procedure, following the conventional milliliter scale protocol [49]. Briefly, a DMSO/NaOH slurry was first prepared by dissolving 300 µL 50 wt % aqueous solution sodium hydroxide (Acros Organics # 25986025) in 600 µL methanol (Fisher Scientific # AF544). This was shaken with 8.0 mL DMSO and centrifuged (3000 x*g*, 3 min, RT). The mixture was decanted, 8.0 mL fresh DMSO was added, mixed well, and centrifuged again. This was repeated three more times to yield a translucent NaOH pellet that was dissolved in 3 mL DMSO and used immediately. Prior to methylation, the dried, reduced glycans were dissolved in 200 µL DMSO and pre-incubated for 10 min at 37°C followed by 20 min shaking at RT. Then 350 µL freshly vortexed NaOH/DMSO slurry was added, followed by an additional 5 min of mixing. To this mixture, 100 µL methyl iodide (Sigma Aldrich # 289566) was added and the sample was vigorously mixed (10 min, RT). The reaction mixture was mixed with 1 mL water and nitrogen gas was bubbled through the solution to remove excess methyl iodide. The permethylated glycans were extracted into 2 mL dichloromethane (Sigma Aldrich # D65100), washed three times with 750 µL water, and dried under a stream of nitrogen.

### LC-MS/MS instrument method and analysis

Permethylated glycan samples were dissolved in 20% MeCN (aq.) prior to LC-MS/MS analysis and stored at -20 °C. Chromatography was performed on an Ultimate 3000 UHPLC (Thermo Scientific) equipped with a reversed phase Waters Acquity Peptide BEH C18 column (150 mm x 2.1 mm inner diameter, 130 Å particle size). Column temperature was maintained at 50-55 °C. Mobile phase A consisted of water (18.2 MΩ) with 0.1% formic acid (Thermo Scientific #28905), and mobile phase B was MeCN (Fisher #A994) with 0.1% formic acid. Flow rate was maintained at 100 or 120 µL/min. The elution gradient was from 20% buffer B to 60% buffer B over span of at least 90 minutes, increased up to 95% B over the final 15 minutes. The UHPLC was interfaced with an LTQ XL ETD Hybrid Ion Trap-Orbitrap ESI mass spectrometer (Thermo Scientific). The mass spectrometer was operated in positive ion mode in the mass range of 700 m/z – 2000 m/z with a spray voltage of 3.5 kV. MS^2^ data was collected in data-dependent acquisition mode, with the top four most abundant ions with signal intensity over 10000 counts selected from the full MS^1^ scan for collision induced dissociation (CID). Calibration was performed regularly (Thermo LTQ ESI Positive Ion Calibration Solution, #88322) to ensure accuracy of 10 ppm or less. Data was processed with XCalibur 2.1 (Thermo Scientific). Relative proportions of glycans were calculated using the area of the extracted ion chromatogram (EIC) for all adducts and charge states for a particular glycan or by integrating the total ion count chromatographic peak (for basigin immunoprecipitation experiments). Glycan compositions and structures were identified using Simglycan [50] and GlycoWorkbench [51],

## Supplemental Information

Supporting information includes supplemental experimental procedures, figures, and tables. This material is available online at XXX. Proteomics datasets are deposited in XXX.

## Author Contributions

E.J., and M. L. H. designed research; E.J., T. O., W. L., R. H., J. R. H. and M. L. H. performed research; E.J., T. O., W. L., C. G. P., and M. L. H. analyzed data; and E.J. and M.L.H. wrote the paper.

## Acknowledgements

E. J. is supported by a Skaggs Graduate Fellowship, enabled by the Henry and Jennifer Luttrell Foundation. M. L. H. is supported by the NIH K99/R00 Pathway to Independence Award (R00HD0292-03). The Huang Lab is supported by startup funds from Scripps Research and the Joe W. and Dorothy Dorsett Brown Foundation.

## Notes

### Competing Interest Statement

The authors have declared no competing interest.

## References

1. Di Lella, S., et al., When galectins recognize glycans: from biochemistry to physiology and back again. Biochemistry, 2011. 50(37): p. 7842–57.

2. Henderson, N.C., et al., Galectin-3 expression and secretion links macrophages to the promotion of renal fibrosis. Am J Pathol, 2008. 172(2): p. 288–98.

3. Henderson, N.C.M. A. C., et al., Galectin-3 regulates myofibroblast activation and hepatic fibrosis. Proc Natl Acad Sci U S A, 2006. 103(13): p. 5060–6065.

4. Rillahan, C.D. and J.C. Paulson, Glycan microarrays for decoding the glycome. Annu Rev Biochem, 2011. 80: p. 797–823.

5. Cederfur, C., et al., Glycoproteomic identification of galectin-3 and -8 ligands in bronchoalveolar lavage of mild asthmatics and healthy subjects. Biochim Biophys Acta, 2012. 1820(9): p. 1429–36.

6. Priglinger, C.S., et al., Galectin-3 induces clustering of CD147 and integrin-beta1 transmembrane glycoprotein receptors on the RPE cell surface. PLoS One, 2013. 8(7): p. e70011.

7. Wang, M., et al., Quantitative proteomics reveal the anti-tumour mechanism of the carbohydrate recognition domain of Galectin-3 in Hepatocellular carcinoma. Sci Rep, 2017. 7(1): p. 5189.

8. Dange, M.C., et al., Mass spectrometry based identification of galectin-3 interacting proteins potentially involved in lung melanoma metastasis. Mol Biosyst, 2017. 13(11): p. 2303–2309.

9. Lakshminarayan, R., et al., Galectin-3 drives glycosphingolipid-dependent biogenesis of clathrin-independent carriers. Nat Cell Biol, 2014. 16(6): p. 595–606.

10. Obermann, J., et al., Proteome-wide Identification of Glycosylation-dependent Interactors of Galectin-1 and Galectin-3 on Mesenchymal Retinal Pigment Epithelial (RPE) Cells. Mol Cell Proteomics, 2017. 16(8): p. 1528–1546.

11. Wang, L., et al., Cross-platform comparison of glycan microarray formats. Glycobiology, 2014. 24(6): p. 507–17.

12. Briard, J.G., et al., Cell-based glycan arrays for probing glycan-glycan binding protein interactions. Nat Commun, 2018. 9(1): p. 880.

13. Ramya, T.N., et al., In situ trans ligands of CD22 identified by glycan-protein photocross-linking-enabled proteomics. Mol Cell Proteomics, 2010. 9(6): p. 1339–51.

14. Malaker, S.A., et al., The mucin-selective protease StcE enables molecular and functional analysis of human cancer-associated mucins. Proc Natl Acad Sci U S A, 2019. 116(15): p. 7278–7287.

15. Trinkle-Mulcahy, L., Recent advances in proximity-based labeling methods for interactome mapping. F1000Res, 2019. 8.

16. Lam, S.S., et al., Directed evolution of APEX2 for electron microscopy and proximity labeling. Nat Methods, 2015. 12(1): p. 51–4.

17. Hung, V., et al., Spatially resolved proteomic mapping in living cells with the engineered peroxidase APEX2. Nat Protoc, 2016. 11(3): p. 456–75.

18. Lepur, A., et al., Ligand induced galectin-3 protein self-association. J Biol Chem, 2012. 287(26): p. 21751–6.

19. Nieminen, J., et al., Visualization of galectin-3 oligomerization on the surface of neutrophils and endothelial cells using fluorescence resonance energy transfer. J Biol Chem, 2007. 282(2): p. 1374–83.

20. Rhee, H.W., et al., Proteomic mapping of mitochondria in living cells via spatially restricted enzymatic tagging. Science, 2013. 339(6125): p. 1328–1331.

21. Bocker, S. and L. Elling, Binding characteristics of galectin-3 fusion proteins. Glycobiology, 2017. 27(5): p. 457–468.

22. Salomonsson, E., et al., Mutational tuning of galectin-3 specificity and biological function. J Biol Chem, 2010. 285(45): p. 35079–91.

23. Ting, L., et al., MS3 eliminates ratio distortion in isobaric multiplexed quantitative proteomics. Nat Methods, 2011. 8(11): p. 937–40.

24. McAlister, G.C., et al., MultiNotch MS3 enables accurate, sensitive, and multiplexed detection of differential expression across cancer cell line proteomes. Anal Chem, 2014. 86(14): p. 7150–8.

25. UniProt, C., UniProt: a worldwide hub of protein knowledge. Nucleic Acids Res, 2019. 47(D1): p. D506–D515.

26. Delaine, T., et al., Galectin-3-Binding Glycomimetics that Strongly Reduce Bleomycin-Induced Lung Fibrosis and Modulate Intracellular Glycan Recognition. Chembiochem, 2016. 17(18): p. 1759–70.

27. Fritsch, K., et al., Galectin-3 interacts with components of the nuclear ribonucleoprotein complex. BMC Cancer, 2016. 16: p. 502.

28. Haudek, K.C., et al., Dynamics of galectin-3 in the nucleus and cytoplasm. Biochim Biophys Acta, 2010. 1800(2): p. 181–9.

29. Mori, Y., et al., Binding of Galectin-3, a beta-Galactoside-binding Lectin, to MUC1 Protein Enhances Phosphorylation of Extracellular Signal-regulated Kinase 1/2 (ERK1/2) and Akt, Promoting Tumor Cell Malignancy. J Biol Chem, 2015. 290(43): p. 26125–40.

30. Talaga, M.L., et al., Multitasking Human Lectin Galectin-3 Interacts with Sulfated Glycosaminoglycans and Chondroitin Sulfate Proteoglycans. Biochemistry, 2016. 55(32): p. 4541–51.

31. Nielsen, M.I., et al., Galectin binding to cells and glycoproteins with genetically modified glycosylation reveals galectin-glycan specificities in a natural context. J Biol Chem, 2018. 293(52): p. 20249–20262.

32. Patnaik, S.K., et al., Complex N-glycans are the major ligands for galectin-1, -3, and -8 on Chinese hamster ovary cells. Glycobiology, 2006. 16(4): p. 305–17.

33. Song, X., et al., Novel fluorescent glycan microarray strategy reveals ligands for galectins. Chem Biol, 2009. 16(1): p. 36–47.

34. Zhou, S., K.M. Wooding, and Y. Mechref, Analysis of Permethylated Glycan by Liquid Chromatography (LC) and Mass Spectrometry (MS). Methods Mol Biol, 2017. 1503: p. 83–96.

35. Chang, L., et al., Identification of Siglec Ligands Using a Proximity Labeling Method. J Proteome Res, 2017. 16(10): p. 3929–3941.

36. Hopkins, C., et al., Chimeric molecules employing horseradish peroxidase as reporter enzyme for protein localization in the electron microscope. Methods Enzymol, 2000. 327: p. 35–45.

37. Charrin, S., et al., Tetraspanins at a glance. J Cell Sci, 2014. 127(Pt 17): p. 3641–8.

38. Stipp, C.S., T.V. Kolesnikova, and M.E. Hemler, EWI-2 is a major CD9 and CD81 partner and member of a novel Ig protein subfamily. J Biol Chem, 2001. 276(44): p. 40545–54.

39. Liang, Y., et al., Complex N-linked glycans serve as a determinant for exosome/microvesicle cargo recruitment. J Biol Chem, 2014. 289(47): p. 32526–37.

40. Wu, M.H., et al., Glycosylation-dependent galectin-1/neuropilin-1 interactions promote liver fibrosis through activation of TGF-beta- and PDGF-like signals in hepatic stellate cells. Sci Rep, 2017. 7(1): p. 11006.

41. Qin, Y., et al., Alteration of protein glycosylation in human hepatic stellate cells activated with transforming growth factor-beta1. J Proteomics, 2012. 75(13): p. 4114–23.

42. Huang, W., et al., Modulation of CD147-induced matrix metalloproteinase activity: role of CD147 N-glycosylation. Biochem J, 2013. 449(2): p. 437–48.

43. Yang, L., et al., Targeted identification of metastasis-associated cell-surface sialoglycoproteins in prostate cancer. Mol Cell Proteomics, 2011. 10(6): p. M110 007294.

44. Liao, C.G., et al., Characterization of basigin isoforms and the inhibitory function of basigin-3 in human hepatocellular carcinoma proliferation and invasion. Mol Cell Biol, 2011. 31(13): p. 2591–604.

45. Ma, T.Y., et al., Cluster of differentiation 147 is a key molecule during hepatocellular carcinoma cell-hepatic stellate cell cross-talk in the rat liver. Molecular Medicine Reports, 2015. 12(1): p. 111–118.

46. Paek, J., et al., Multidimensional Tracking of GPCR Signaling via Peroxidase-Catalyzed Proximity Labeling. Cell, 2017. 169(2): p. 338–349 e11.

47. Xu, L., et al., Human hepatic stellate cell lines, LX-1 and LX-2: new tools for analysis of hepatic fibrosis. Gut, 2005. 54(1): p. 142–51.

48. Hamouda, H., et al., Rapid analysis of cell surface N-glycosylation from living cells using mass spectrometry. J Proteome Res, 2014. 13(12): p. 6144–51.

49. Shajahan, A., et al., High-Throughput Automated Micro-permethylation for Glycan Structure Analysis. Anal Chem, 2019. 91(2): p. 1237–1240.

50. Apte, A. and N.S. Meitei, Bioinformatics in glycomics: glycan characterization with mass spectrometric data using SimGlycan. Methods Mol Biol, 2010. 600: p. 269–81.

51. Ceroni, A., et al., GlycoWorkbench: A Tool for the Computer-Assisted Annotation of Mass Spectra of Glycans. Journal of Proteome Research, 2008. 7(4): p. 1650–1659.

