## Supporting Information for "Mapping glycan-mediated galectin-3 interactions by live cell proximity labeling"

**CONTENTS:**

**Figure S1.** Additional design and characterization data for PX-Gal3 and PX-Gal3$_{\Delta116}$ fusion proteins.

**Figure S2.** LCMS analysis of asialofetuin.

**Figure S3.** Additional data on the use of the proximity labeling method in live cells.

**Figure S4.** Additional fluorescence microscopy images for PX-Gal3$_{tm}$.

**Figure S5.** Full Western blot of Figure 2E.

**Figure S6.** Additional data from quantitative proteomics experiments collected from exogenous incubations.

**Figure S7.** Additional data collected from transfection experiments.

**Figure S8.** Western blots to show endogenous expression of proteins in LX-2 cells.

**Table S1.** Commercial recombinant human proteins used for ELISA assay.

**Table S2.** Curve fitting analyses for ELISA data in Figure 4B.

**Figure S9.** Full comparison of glycomics data from cell surfaces, PX-Gal3 enriched glycoproteins, and immunoprecipitated basigin.

**Figure S10.** Additional glycomics data collected for LX-2 cell surfaces.

**Figure S11.** Cell surface staining of LX-2 cells using plant lectins.

**Figure S12.** Additional glycomics data collected for PX-Gal3 enriched N-glycans.

**Figure S13.** Comparison of PX-Gal3 enriched and cell surface N-glycans.

**Figure S14.** Full SDS-PAGE and Western of immunoprecipitated basigin.

**Figure S15.** Additional glycomics data collected for basigin-IP sample.

**Figure S16.** Glycosite analysis of basigin-Fc recombinant protein.

**APPENDIX:**

**Table S1.** Proteomics data for proximity labeling of PX-Gal3 fusion proteins.

A

PX-Gal3 (1553 bp, 62% GC)
His$_6$-APEX2-linker-Gal-3

```
1     GAATTCATGC ATCACCACCA CCACCACGGT AAAAGCTACC CGACCGTGAG CGCGGACTAT
61    CAGGATGCGG TTGAAAAAGC GAAGAAAAAG CTGCGTGGTT TTATCGCGGA GAAGCGTTGC
121   GCGCCGCTGA TGCTGCGTCT GGCGTTCCAC AGCGCGGGTA CCTTTGACAA AGGTACCAAG
181   ACCGGTGGCC CGTTCGGCAC CATTAAACAT CCGGCGGAAC TGGCGCACAG CGCGAACAAC
241   GGTCTGGACA TCGCGGTGCG TCTGCTGGAA CCGCTGAAGG CGGAGTTCCC GATTCTGAGC
301   TACGCGGATT TTTATCAGCT GGCGGGCGTG GTTGCGGTTG AGGTGACCGG TGGCCCGAAA
361   GTTCCGTTCC ACCCGGGTCG TGAGGACAAA CCGGAGCCGC CGCCGGAGGG CCGTCTGCCG
421   GATCCGACCA AAGGTAGCGA CCACCTGCGT GATGTGTTTG GCAAGGCGAT GGGTCTGACC
481   GACCAAGATA TCGTTGCGCT GAGCGGTGGC CACACCATTG GTGCGGCGCA CAAGGAACGT
541   AGCGGTTTCG AGGGCCCGTG GACCAGCAAC CCGCTGATCT TCGACAACAG CTACTTTACC
601   GAACTGCTGA GCGGCGAGAA AGAAGGTCTG CTGCAGCTGC CGAGCGACAA GGCGCTGCTG
661   AGCGATCCGG TGTTCCGTCC GCTGGTTGAC AAATACGCGG CGGACGAGGA TGCGTTCTTT
721   GCGGATTATG CGGAAGCGCA CCAAAAACTG AGCGAGCTGG GTTTTGCGGA TGCGGGTAGC
781   GGTGGCGGTG GCAGCGCGGA CAACTTTAGC CTGCACGATG CGCTGAGCGG TAGCGGTAAC
841   CCGAACCCGC AGGGTTGGCC GGGTGCGTGG GGTAACCAAC CGGCGGGTGC GGGTGGCTAC
901   CCGGGTGCGA GCTATCCGGG TGCGTATCCG GGTCAGGCTC CGCCGGGTGC GTACCCGGGC
961   CAAGCTCCGC CGGGTGCTTA TCCTGGTGCG CCGGGCGCGT ACCCGGGTGC GCCGGCGCCG
1021  GGCGTGTACC CGGGTCCGCC GAGCGGTCCG GGCGCGTATC CGAGCAGCGG CCAGCCGAGC
1081  GCGACCGGTG CGTACCCGGC GACCGGCCCG TATGGTGCGC CGGCGGGTCC GCTGATTGTT
1141  CCGTATAACC TGCCGCTGCC GGGTGGCGTG GTTCCGCGTA TGCTGATCAC CATTCTGGGC
1201  ACCGTGAAAC CGAACGCGAA CCGTATCGCG CTGGACTTCC AACGTGGTAA CGATGTTGCG
1261  TTCCACTTTA ACCCGCGTTT TAACGAAAAC AACCGTCGTG TGATTGTTTG CAACACCAAG
1321  CTGGACAACA ACTGGGGCCG TGAGGAACGT CAGAGCGTGT TCCCGTTTGA AAGCGGTAAA
1381  CCGTTCAAGA TTCAAGTGCT GGTTGAGCCG GACCACTTTA AAGTGGCGGT TAACGATGCG
1441  CACCTGCTGC AATACAACCA CCGTGTTAAA AAGCTGAACG AGATCAGCAA GCTGGGCATC
1501  AGCGGTGACA TTGATCTGAC CAGCGCGAGC TATACCATGA TTTAAGCGGC CGC
```

PX-Gal3$_{\Delta 116}$ (1226 bp)
His6-APEX2-(Δ1-116)Gal-3

```
1     GAATTCATGC ATCACCACCA CCACCACGGT AAAAGCTACC CGACCGTGAG CGCGGACTAT
61    CAGGATGCGG TTGAAAAAGC GAAGAAAAAG CTGCGTGGTT TTATCGCGGA GAAGCGTTGC
121   GCGCCGCTGA TGCTGCGTCT GGCGTTCCAC AGCGCGGGTA CCTTTGACAA AGGTACCAAG
181   ACCGGTGGCC CGTTCGGCAC CATTAAACAT CCGGCGGAAC TGGCGCACAG CGCGAACAAC
241   GGTCTGGACA TCGCGGTGCG TCTGCTGGAA CCGCTGAAGG CGGAGTTCCC GATTCTGAGC
301   TACGCGGATT TTTATCAGCT GGCGGGCGTG GTTGCGGTTG AGGTGACCGG TGGCCCGAAA
361   GTTCCGTTCC ACCCGGGTCG TGAGGACAAA CCGGAGCCGC CGCCGGAGGG CCGTCTGCCG
421   GATCCGACCA AAGGTAGCGA CCACCTGCGT GATGTGTTTG GCAAGGCGAT GGGTCTGACC
481   GACCAAGATA TCGTTGCGCT GAGCGGTGGC CACACCATTG GTGCGGCGCA CAAGGAACGT
541   AGCGGTTTCG AGGGCCCGTG GACCAGCAAC CCGCTGATCT TCGACAACAG CTACTTTACC
601   GAACTGCTGA GCGGCGAGAA AGAAGGTCTG CTGCAGCTGC CGAGCGACAA GGCGCTGCTG
661   AGCGATCCGG TGTTCCGTCC GCTGGTTGAC AAATACGCGG CGGACGAGGA TGCGTTCTTT
721   GCGGATTATG CGGAAGCGCA CCAAAAACTG AGCGAGCTGG GTTTTGCGGA TGCGGGTAGC
781   GGTGGCGGTG GCAGCCCGTA TAACCTGCCG CTGCCGGGTG GCGTGGTTCC GCGTATGCTG
841   ATCACCATTC TGGGCACCGT GAAACCGAAC GCGAACCGTA TCGCGCTGGA CTTCCAACGT
901   GGTAACGATG TTGCGTTCCA CTTTAACCCG CGTTTTAACG AAAACAACCG TCGTGTGATT
961   GTTTGCAACA CCAAGCTGGA CAACAACTGG GGCCGTGAGG AACGTCAGAG CGTGTTCCCG
1021  TTTGAAAGCG GTAAACCGTT CAAGATTCAA GTGCTGGTTG AGCCGGACCA CTTTAAAGTG
1081  GCGGTTAACG ATGCGCACCT GCTGCAATAC AACCACCGTG TTAAAAAGCT GAACGAGATC
1141  AGCAAGCTGG GCATCAGCGG TGACATTGAT CTGACCAGCG CGAGCTATAC CATGATTTAA
1201  GCGGCCGC
```

B

PX-Gal3
His$_6$-APEX2-linker-Gal-3
512 aa; 51 kDa

MHHHHHHGKSYPTVSADYQDAVEKAKKKLRGFIAEKRCAPLMLRLAFHSAGTFDKGTKTGGPFGTIKHPAELAHSANNGLDIAVRLLEPLKAEF
PILSYADFYQLAGVVAVEVTGGPKVPFHPGREDKPEPPPEGRLPDPTKGSDHLRDVFGKAMGLTDQDIVALSGGHTIGAAHKERSGFEGPWTSNPLIFDNS
YFTELLSGEKEGLLQLPSDKALLSDPVFRPLVDKYAADEDAFFADYAEAHQKLSELGFADAGSGGGGSADNFSLHDALSGSGNPNPQGWPGAWGNQPAG
AGGYPGASYPGAYPGQAPPGAYPGQAPPGAYPGAPGAYPGAPAPGVYPGPPSGPGAYPSSGQPSATGAYPATGPYGAPAGPLIVPYNLPLPGGVVPRML
ITILGTVKPNANRIALDFQRGNDVAFHFNPRFNENNRRVIVCNTKLDNNWGREERQSVFPFESGKPFKIQVLVEPDHFKVAVNDAHLLQYNHRVKKLNEIS
KLGISGDIDLTSASYTMI*

PX-Gal3$_{\Delta 116}$
His6-APEX2-linker-(Δ1-116)Gal-3
403 aa; 40 kDa

MHHHHHHMGKSYPTVSADYQDAVEKAKKKLRGFIAEKRCAPLMLRLAFHSAGTFDKGTKTGGPFGTIKHPAELAHSANNGLDI
AVRLLEPLKAEFPILSYADFYQLAGVVAVEVTGGPKVPFHPGREDKPEPPPEGRLPDPTKGSDHLRDVFGKAMGLTDQDIVALSGGHTIGA
AHKERSGFEGPWTSNPLIFDNSYFTELLSGEKEGLLQLPSDKALLSDPVFRPLVDKYAADEDAFFADYAEAHQKLSELGFADAGAPRARARP
RARAYNLPLPGGVVPRMLITILGTVKPNANRIALDFQRGNDVAFHFNPRFNENNRRVIVCNTKLDNNWGREERQSVFPFESGKPFKIQVL
VEPDHFKVAVNDAHLLQYNHRVKKLNEISKLGISGDIDLTSASYTMI*

C

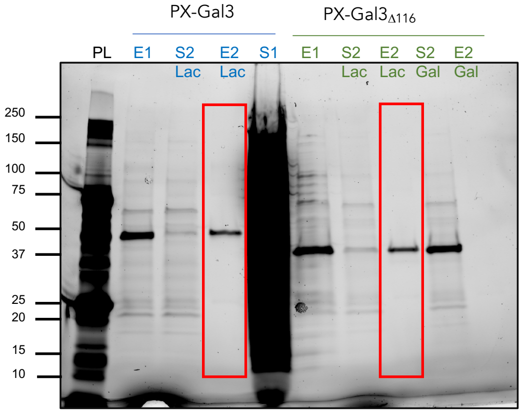


D

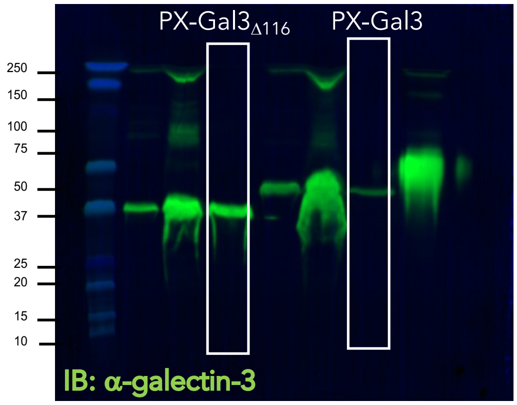


**Figure S1. Expression of APEX2-galectin-3 fusion constructs.** (**A**) DNA sequence of inserted PX-Gal3 and PX-Gal3$_{\Delta 116}$ constructs, including nucleotide sequences used for cloning. **(B)** Predicted translated protein sequences. **(C)** SDS-PAGE analysis of proteins eluted from nickel beads or columns (E1) and further purification with lactose beads (E2). (**D**) Western blotting for galectin-3 shows that the resulting constructs are immuno-positive.

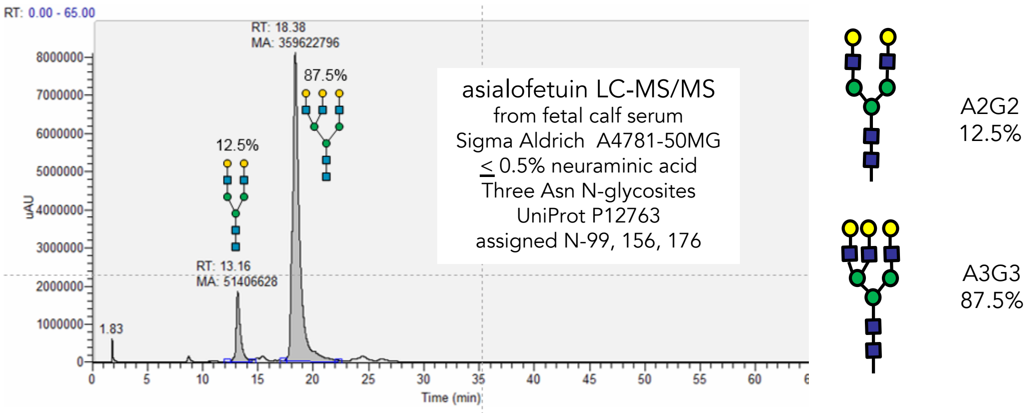


**Figure S2. LCMS analysis of N-glycans present in commercial asialofetuin used in ELISA experiments (Fig. 2B)**. Terminal galactose residues (yellow circle), which are high affinity ligands for galectin-3, are abundant.

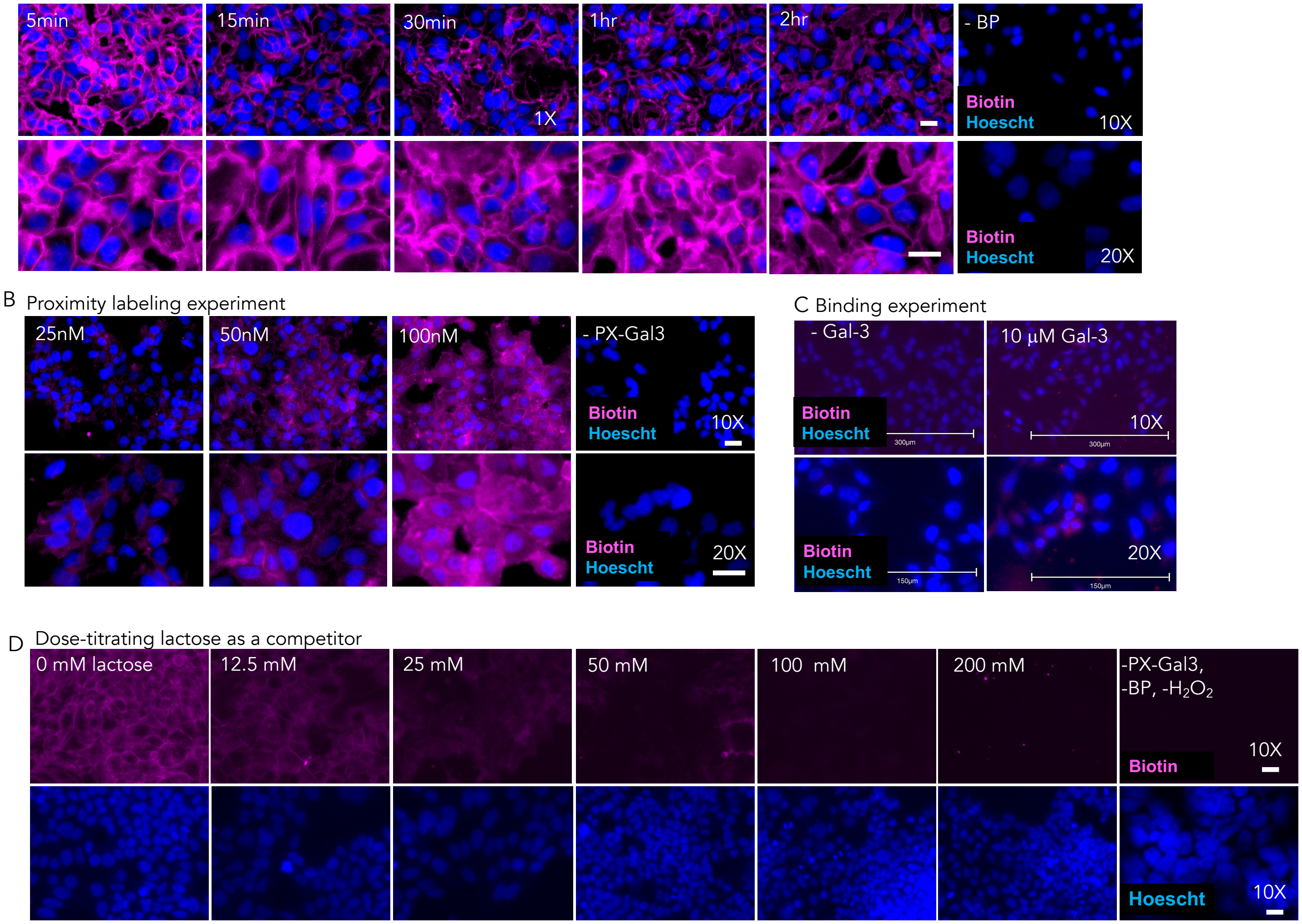


**Figure S3. Fluorescence micrographs to show parameters affecting in situ proximity labeling in live cells.** Cells were stained for biotinylated interactors with Cy5-Streptavidin (purple) and nuclei are stained with Hoescht 33342 (blue). Top and bottom panels show images at 10x and 20x magnification, respectively. Scale bars: 30 μm **(A)** Five-minute incubation of LX-2 cells with PX-Gal3 (100 nM) is sufficient to achieve robust labeling. **(B)** There is a dose-dependent increase in labeling of interactors PX-Gal3 following 30 mins of protein incubation (37°C). **(C)** Detection of the binding of galectin-3 (10 μM) to LX-2s requires significantly higher protein concentrations and incubation periods (overnight, 4°C). **(D)** There is significant loss of fluorescence following co-incubation of PX-Gal-3 (full, 100 nM, 30 min) with lactose, even at 12.5 mM. Full competition was observed at 100 mM lactose concentrations.

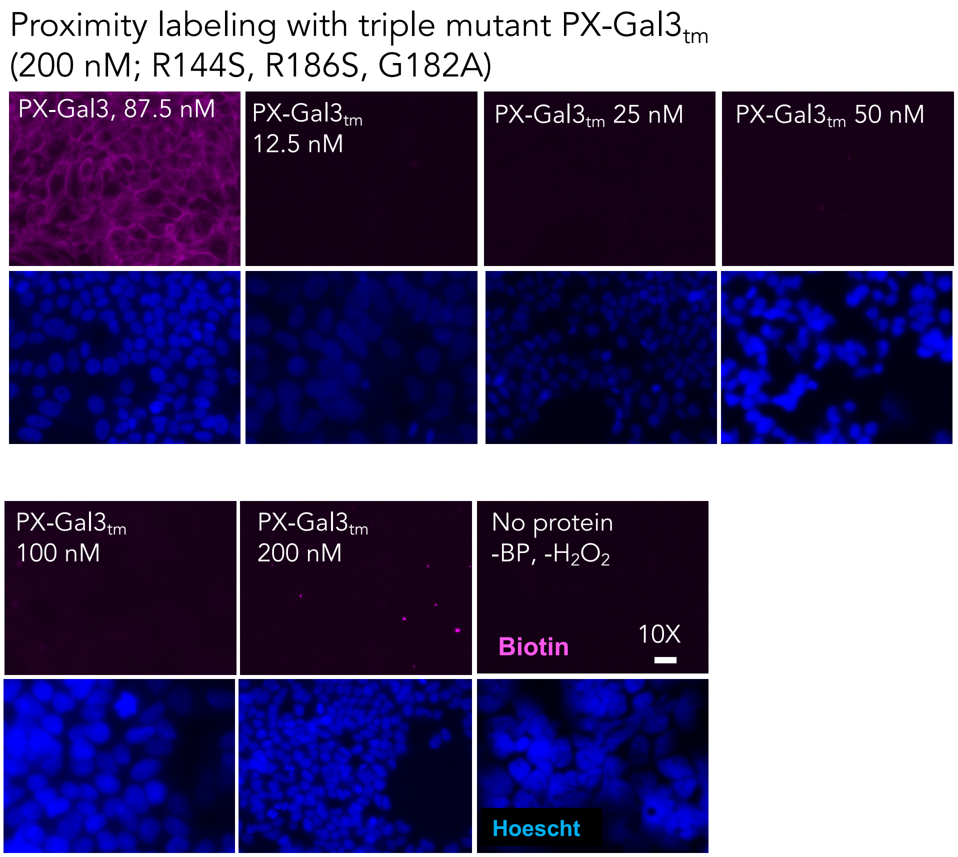


**Figure S4.** A mutant of PX-Gal3 (PX-Gal-$3_{tm}$), where three of the most important residues required for binding glycans in galectin-3 are mutated: R144S, R186S, G182A, show poor proximity labeling, indicating that glycan-mediated interactions are required for binding to LX-2s.

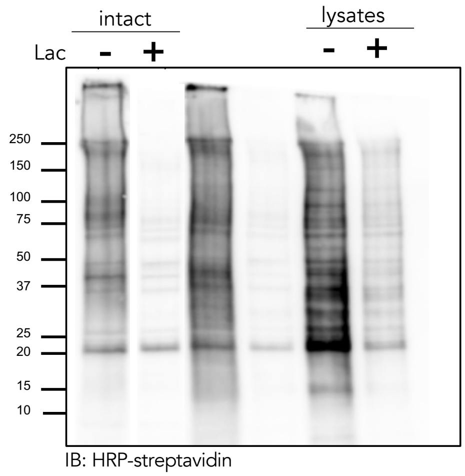


**Figure S5. Full-length Western blot to complement Fig. 2E.** Only a subset of proteins are labeled in intact cells compared to lysates.

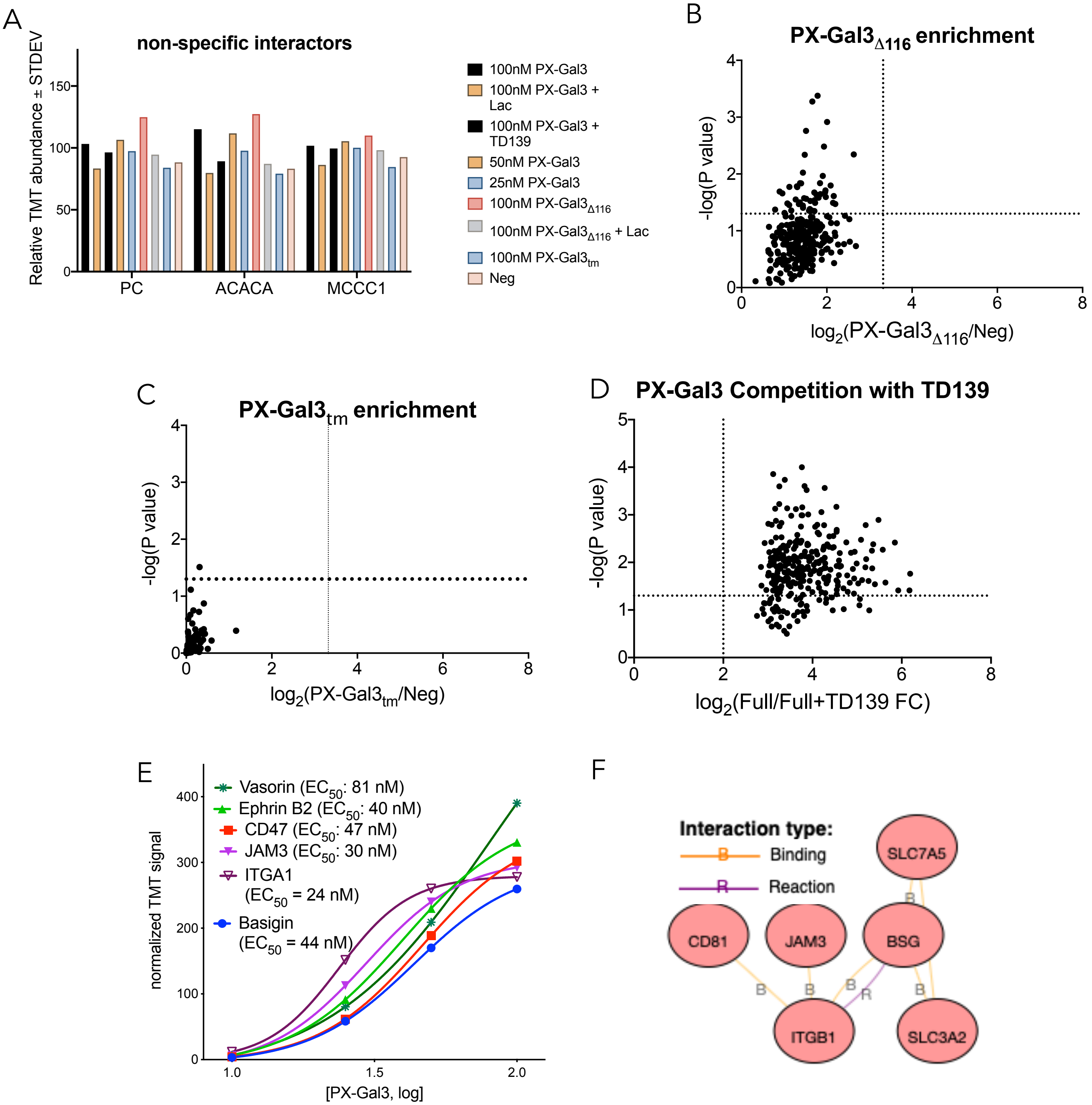


**Figure S6. Additional data from quantitative proteomics analyses collected from exogenous incubations**. **(A)** Non-specific proteins show poor enrichment over negative controls. (**B**) PX-Gal3$_{\Delta 116}$ (100 nM) does not significantly enrich proteins over a negative control (Neg). (**C**) PX-Gal3$_{\tau\mu}$ does not significantly enrich proteins over Neg. (**D**) Interactions of PX-Gal3 are competed by co-incubation with TD139 (15.4 μM). (**E**) Apparent binding affinity values ($EC_{50}$) determined for selected proteins using quantitative MS-based proteomics. (**F**) Integrated Pathway Analysis of binding relationships based on PX-Gal3 enriched proteins.

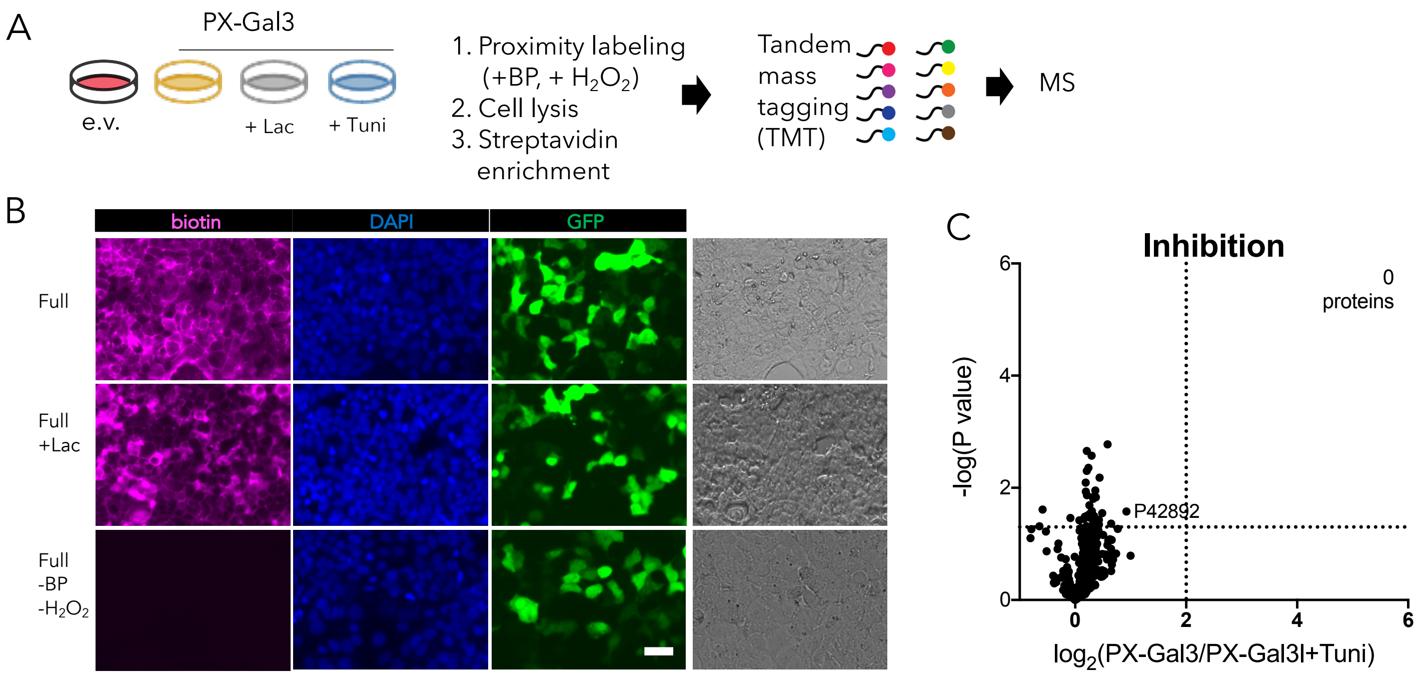


**Figure S7. Additional data collected from transfection experiments. (A)** LX-2 cells were transiently transfected with either empty vector (e.v.) or vector carrying PX-Gal3. Following transfection, cells were additionally treated with excess lactose (100 mM) or tunicamycin (10 μg/mL) for 24 hours, prior to performing proximity labeling. (**B**) In contrast to the exogenous treatments of the fusion proteins, addition of exogenous lactose to cells failed to significantly compete for interactions, most likely due to its intracellular impermeability. Scale bar: 20 μm (**C**) Treatment with tunicamycin, an inhibitor of N-linked glycosylation, did not compete for PX-Gal3 interactions.

| | Full-length protein | Recombinant protein | | | |
|---|---|---|---|---|---|
| | Assigned N-glycosites of full-length protein (UniProt) | Accession # | Cat. # | Predicted MW /Observed MW | Sequence of Recombinant Protein |
| **basigin-2-Fc** | N-44, N-160 of basigin-2 (equivalent to N-160 and 268 of basigin-1) | NP_940991.1 | 10186-H02H | 46.8 kDa/ 58-65 kDa | extracellular domain (Met 1-His 205) of human CD147 basigin-2 precursor was expressed with the fused Fc region of human IgG1 at the C-terminus. *** We confirmed the presence of N-glycans on residues Asn 44 and 152 of basigin-2 and Asn-73 and 299 of the Fc domain*** |
| **CD81** | none | NP_004347.1 | 14244-HNCH | 9.9 kDa/ 10 kDa | (Phe113-Lys201) was expressed with two additional amino acids (Gly&Pro) at the N-terminus. |
| **vasorin** | N-101, 117, 273, 500, 528 | Q6EMK4 | 13854-H08H | 61.1 kDa / 66-76 kDa | (Met1-Pro575) was expressed with a polyhistidine tag at the C-terminus. |
| **ephrin-B1** | N-139 | NP_004420.1 | 10894-H08H | 24.5 kDa/ 38 kDa | extracellular domain (Met 1-Lys 237) was fused with a polyhistidine tag at the C-terminus. |
| **neuroplastin** | N-171, 197, 229, 284, 296, 317 | NP_059429.1 | 15881-H08H | 23.2 kDa | (Met1-Leu221) was expressed with a polyhistidine tag at the C-terminus. *** missing 4 glycosylation sites*** |
| **CD9** | N-52, 53 | NP_001760.1 | 11029-H08H | 10 kDa / 11 kDa | second extracellular domain (Ser 112-Ile 195) of human CD9 was fused with a polyhistidine tag at the C-terminus and a signal peptide at the N-terminus. *** missing both glycosylation sites *** |
| **CD47** | N-23, 34, 50, 73, 111, 206 | NP_942088.1 | 12283-HCCH | 14.5 kDa | (Met1-Pro139) was expressed with six amino acids (LEVLFQ) at the C-terminus. *** missing last Asn-206 glycosylation site *** |

**Table S1. Commercially purchased recombinant proteins used for ELISAs.** Note that these proteins may not reflect endogenous protein sequences and glycosylation states found in LX-2 cells.

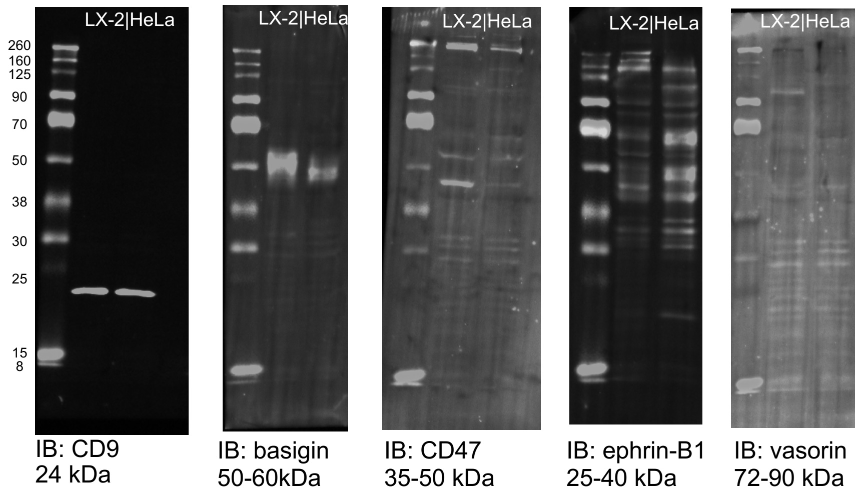


**Figure S8. Western blots showing endogenous expression of various proteins in LX-2 cells.** 10 μg each of cell lysates from LX-2 HSCs or from HeLa (human cervical cancer cells).

| | rhBasigin | rhCD81 | rhvasorin | rhEphrin-B1 | rh-Neuroplastin | rh-CD9 | rh-CD47 |
|---|---|---|---|---|---|---|---|
| log(agonist) vs. response (three parameters) | | | | | | | |
| Best-fit values | | | | | | | |
| Bottom | 0.03046 | 0.007451 | 0.007233 | -0.007259 | 0.03977 | -0.02879 | 0.02969 |
| Top | 1.131 | 1.097 | 1.115 | 1.134 | 1.191 | 1.208 | 1.192 |
| LogEC50 | 3.377 | 3.048 | 3.217 | 3.114 | 3.619 | 2.851 | 3.613 |
| EC50 | 2385 | 1117 | 1649 | 1302 | 4156 | 709.7 | 4100 |
| Span | 1.101 | 1.089 | 1.107 | 1.141 | 1.151 | 1.237 | 1.162 |
| 95% CI (profile likelihood) | | | | | | | |
| Bottom | 0.004798 to 0.05585 | -0.03674 to 0.05078 | -0.01808 to 0.03228 | -0.06184 to 0.04618 | -0.006713 to 0.08528 | -0.1294 to 0.06803 | 0.01379 to 0.04547 |
| Top | 1.070 to 1.197 | 1.026 to 1.172 | 1.065 to 1.167 | 1.041 to 1.233 | 1.042 to 1.385 | 1.078 to 1.348 | 1.137 to 1.251 |
| LogEC50 | 3.284 to 3.471 | 2.919 to 3.177 | 3.137 to 3.298 | 2.961 to 3.267 | 3.415 to 3.831 | 2.630 to 3.067 | 3.544 to 3.683 |
| EC50 | 1925 to 2959 | 829.6 to 1503 | 1370 to 1986 | 915.0 to 1849 | 2602 to 6775 | 426.7 to 1166 | 3496 to 4818 |
| Goodness of Fit | | | | | | | |
| Degrees of Freedom | 18 | 18 | 18 | 18 | 18 | 18 | 18 |
| R squared | 0.9915 | 0.9821 | 0.9931 | 0.9731 | 0.9673 | 0.9415 | 0.9961 |
| Sum of Squares | 0.02416 | 0.06055 | 0.02197 | 0.09772 | 0.08471 | 0.2838 | 0.01006 |
| Sy.x | 0.03663 | 0.05800 | 0.03493 | 0.07368 | 0.06860 | 0.1256 | 0.02364 |
| Number of points | | | | | | | |
| # of X values | 21 | 21 | 21 | 21 | 21 | 21 | 21 |
| # Y values analyzed | 21 | 21 | 21 | 21 | 21 | 21 | 21 |

**Table S1. Curve fitting analyses of ELISA data in Figure 4B.**

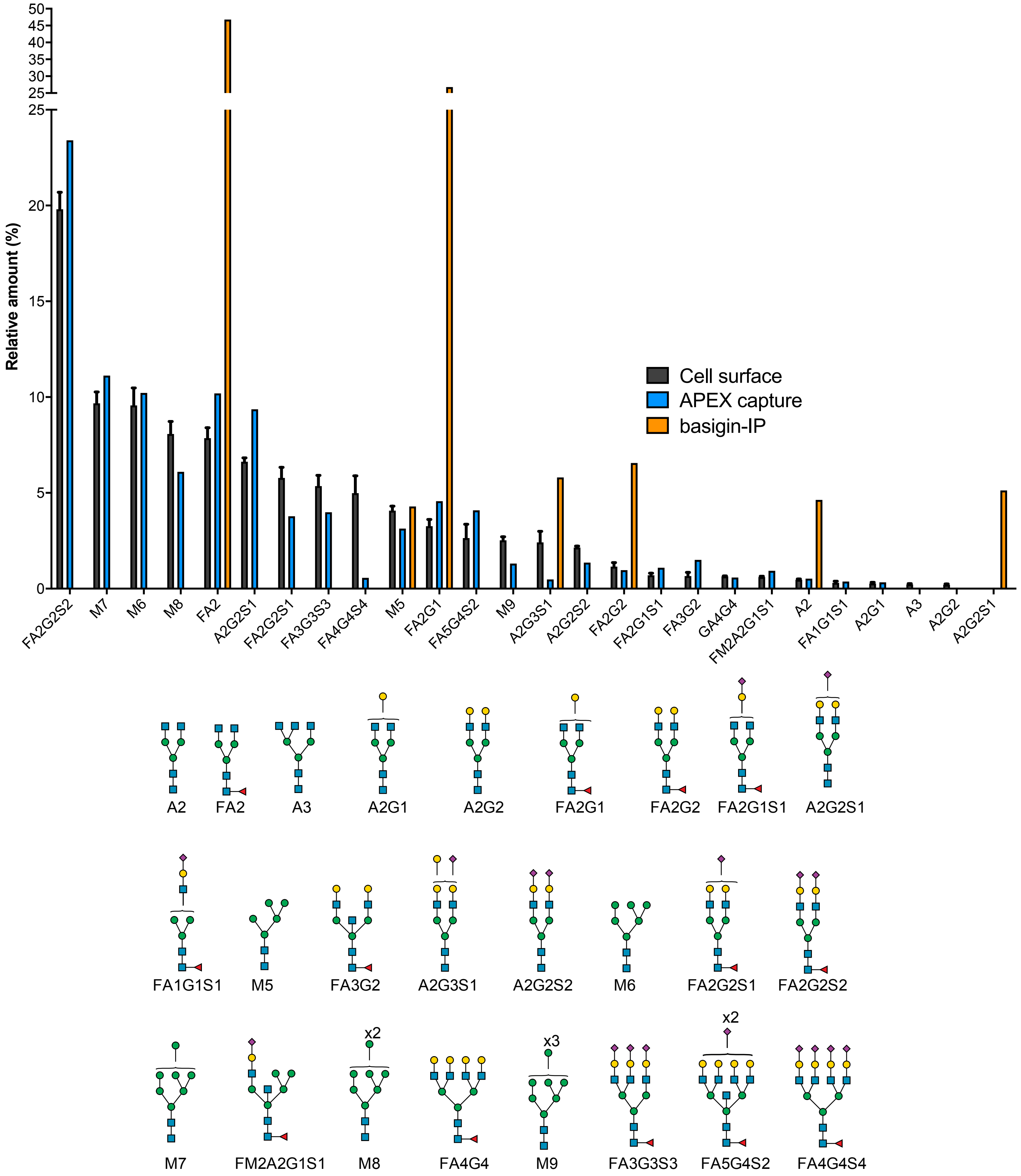


**Figure S9. Full composition analysis of the most abundant N-glycans found on LX-2 cell surfaces, captured by PX-Gal3, or from immunoprecipitated (IP) basigin.** Top panel represents the relative amounts of each N-glycan found in LX-2 cell surfaces, PX-Gal3 enriched samples, or basigin-IP. Bottom panel shows key to N-glycan nomenclature used.

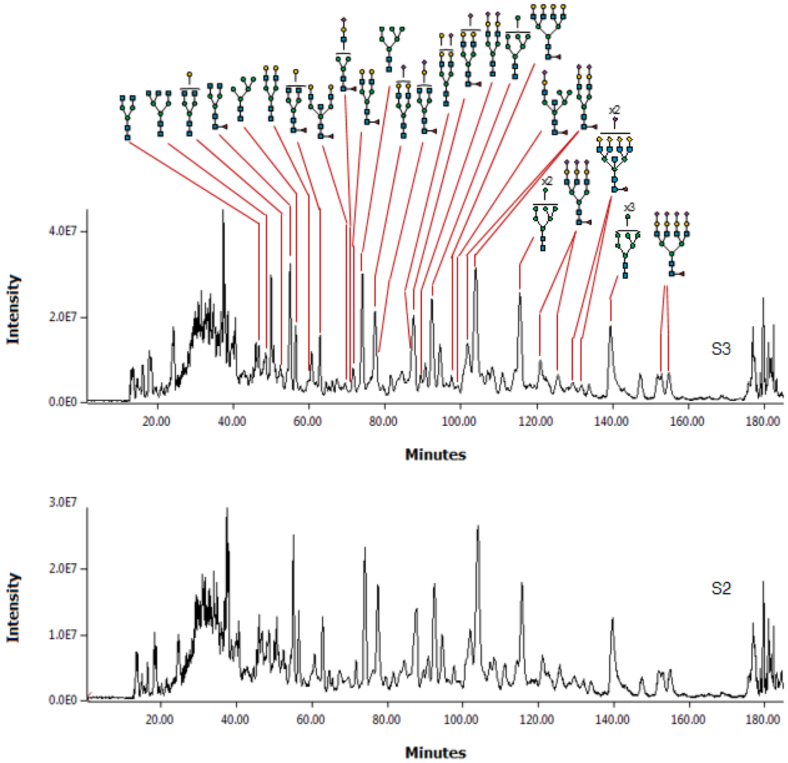


**Figure S10. Glycomics MS analysis of LX-2 cell surfaces**. Total ion chromatograms of permethylated N-glycans from two independent biological replicates (S3, S2).

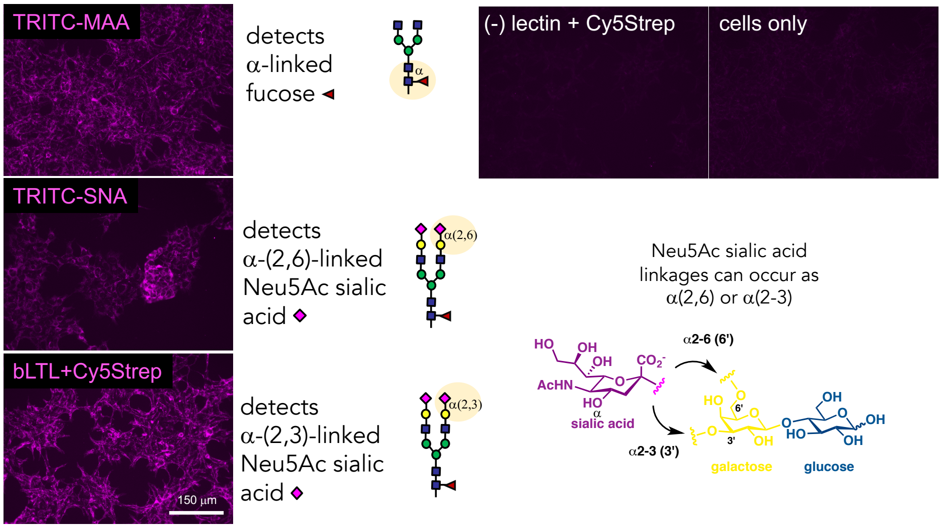


**Figure S11. Lectin staining of LX-2 hepatic stellate cells**. HSCs stain positively for the presence of α-linked fucoses and both α(2,6) and α(2,-3)-linked sialic acid linkages. LX-2 cells were seeded overnight in DMEM+ 10% FBS onto pre-coated (>10 min, RT, extensive washing with DPBS after) poly-L-lysine coated glass slide chambers. Next days, cells were fixed in 4% PFA/PBS (10min, RT) and blocked (2% BSA/PBS + 0.1% TX, 1hr, RT). Lectins (either TRITC-MAA, TRITC-SNA or bLTL; 1:200 in same buffer) were incubated for 1 hr at RT. The bLTL condition was washed and probed with Cy5-streptavidin (1:500 in same buffer). After washing and removal of the chambers, antifade was added and coverslips were attached. Cells were imaged with the appropriate filters.

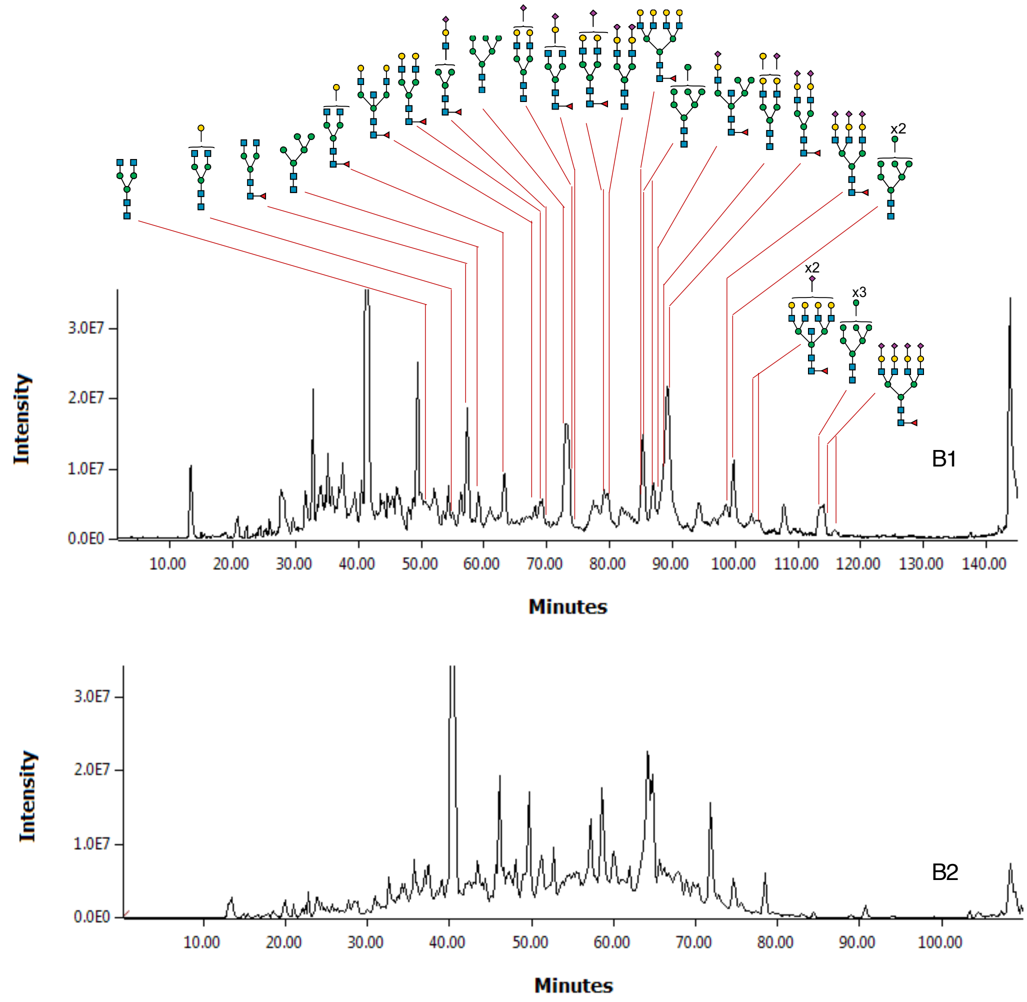


**Figure S12. Additional data for PX-Gal3 enriched glycans.** Total ion chromatograms of permethylated N-glycans from two independent biological replicates (B1, B2). Note that a shorter run time was used in B2.

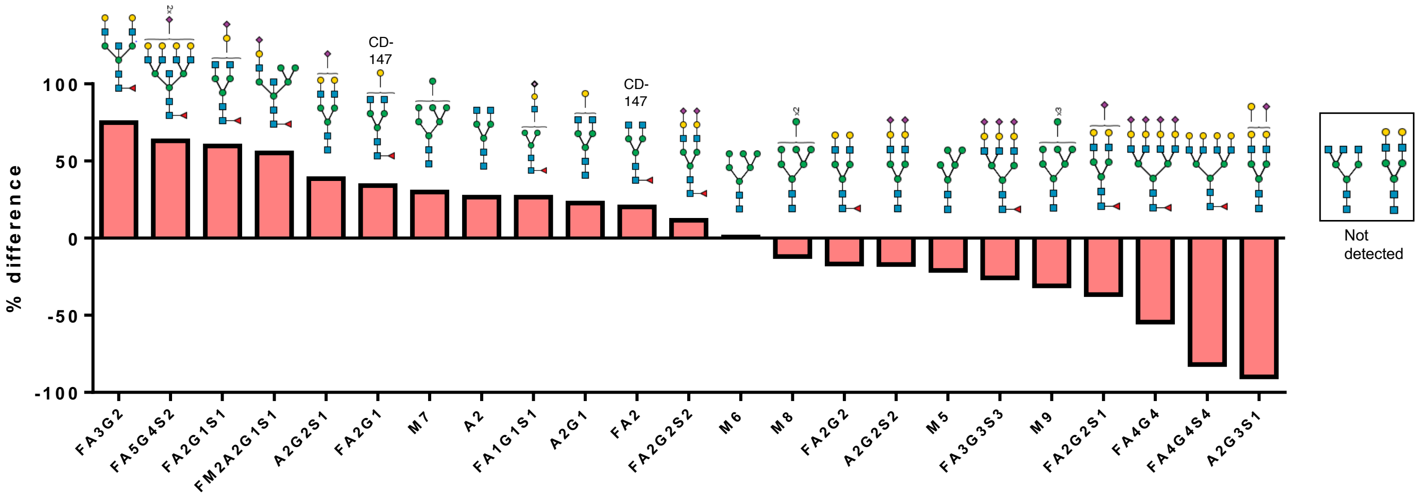

**Figure S13. Full Comparison of PX-Gal3 enriched and cell surface N-glycans.** % difference was calculated as the difference between N-glycan abundance between PX-Gal3 and cell surface divided by the abundance of the cell surface N-glycan.

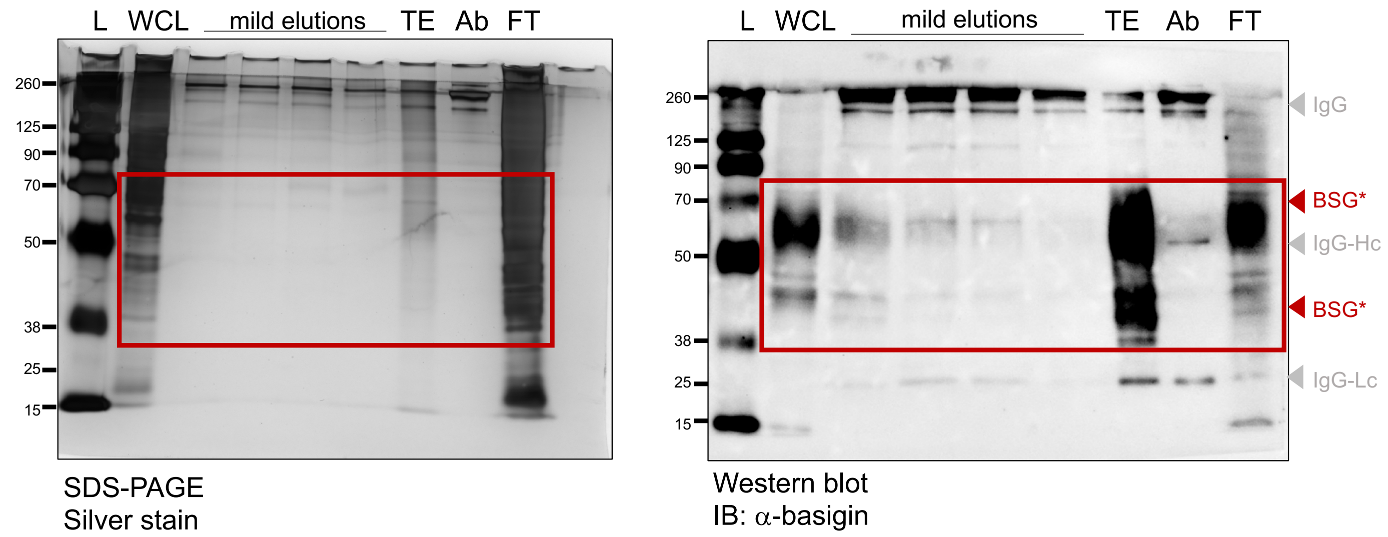


L: ladder
WCL: whole cell lysate
TE: total elution
Ab: antibody
FT: bead flow-through
BSG: basigin
Hc: heavy chain
Lc: light chain

**Figure S14. SDS-PAGE and Western blot analysis of basigin-IP (immunoprecipitated) from LX-2 cells.**

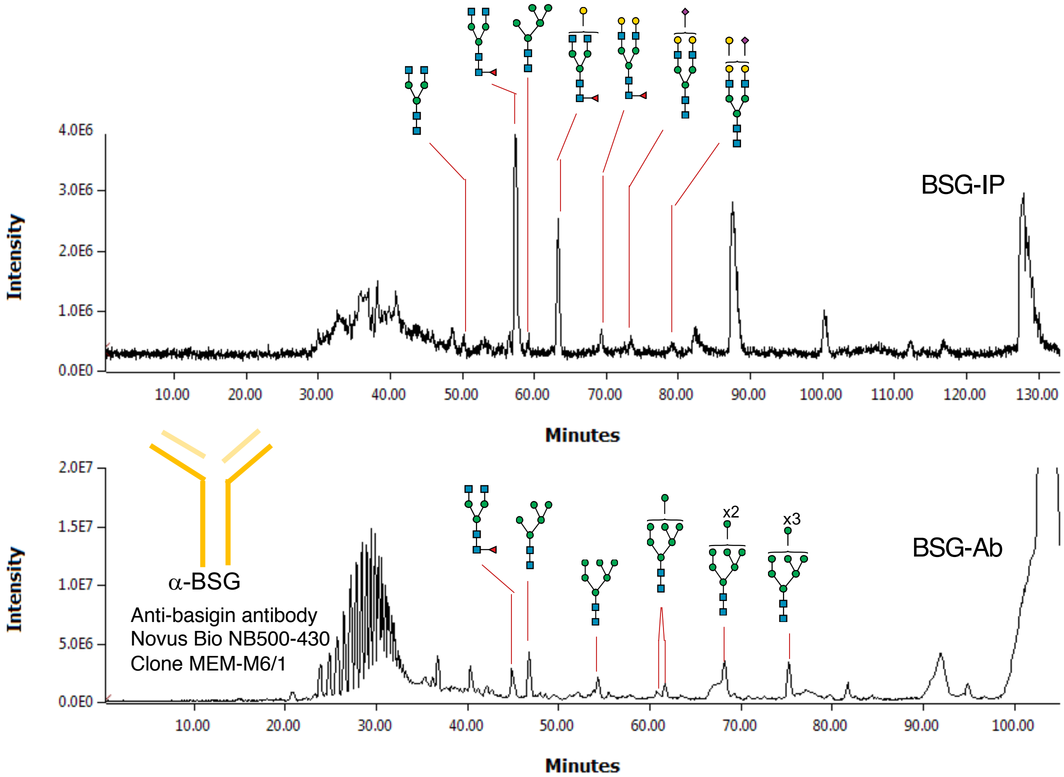


**Figure S15. Additional glycomics data for BSG-IP.** Total ion chromatograms of permethylated N-glycans from a basigin-IP sample. Bottom chromatogram represents the N-glycans from the immunoprecipitating antibody alone. Note that a shorter run time was used in BSG-AB.

### Basigin-2-Fc

Recombinant protein
Sinobio 10186-H02H

#### Basigin-1 vs. Basigin-2 isoform comparison

```
P35613   BASI_HUMAN      1   MAAALFVLLGFALLGTHGASGAAGFVQAPLSQQRWVGGSVELHCEAVGSPVPEIQWWFEG   60
P35613-2 BASI_HUMAN      1   MAAALFVLLGFALLGTHGASG---------------------------------------   21
                             *********************

P35613   BASI_HUMAN     61   QGPNDTCSQLWDGARLDRVHIHATYHQHAASTISIDTLVEEDTGTYECRASNDPDRNHLT  120
P35613-2 BASI_HUMAN     22   ------------------------------------------------------------   21

P35613   BASI_HUMAN    121   RAPRVKWVRAQAVVLVLEPGTVFTTVEDLGSKILLTCSLNDSATEVTGHRWLKGGVVLKE  180
P35613-2 BASI_HUMAN     22   -----------------AAGTVFTTVEDLGSKILLTCSLNDSATEVTGHRWLKGGVVLKE   64
                                              *******************************************

P35613   BASI_HUMAN    181   DALPGQKTEFKVDSDDQWGEYSCVFLPEPMGTANIQLHGPPRVKAVKSSEHINEGETAML  240
P35613-2 BASI_HUMAN     65   DALPGQKTEFKVDSDDQWGEYSCVFLPEPMGTANIQLHGPPRVKAVKSSEHINEGETAML  124
                             ************************************************************

P35613   BASI_HUMAN    241   VCKSESVPPVTDWAWYKITDSEDKALMNGSESRFFVSSSQGRSELHIENLNMEADPGQYR  300
P35613-2 BASI_HUMAN    125   VCKSESVPPVTDWAWYKITDSEDKALMNGSESRFFVSSSQGRSELHIENLNMEADPGQYR  184
                             ************************************************************

P35613   BASI_HUMAN    301   CNGTSSKGSDQAIITLRVRSHLAALWPFLGIVAEVLVLVTIIFIYEKRRKPEDVLDDDDA  360
P35613-2 BASI_HUMAN    185   CNGTSSKGSDQAIITLRVRSHLAALWPFLGIVAEVLVLVTIIFIYEKRRKPEDVLDDDDA  244
                             ************************************************************

P35613   BASI_HUMAN    361   GSAPLKSSGQHQNDKGKNVRQRNSS                                     385
P35613-2 BASI_HUMAN    245   GSAPLKSSGQHQNDKGKNVRQRNSS                                     269
                             *************************
```

▬ Found in recombinant protein

▬ Missing in recombinant protein

#### N-glycosite identification:

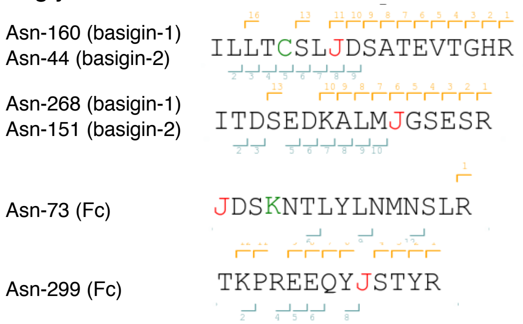


#### N-glycosite mapping:

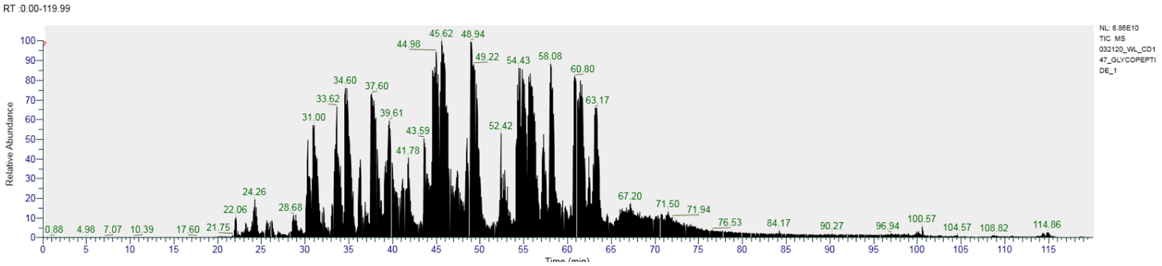


**Figure S16. Analysis of N-linked glycosites in recombinant basigin-Fc.** 20 µg of basigin-Fc (SinoBio # 10186-H02H) was dissolved in 50 µL of 50 mM ammonium bicarbonate buffer. To this, 10 µL of 50 mM dithiothreitol (DTT) was added and incubated at 55°C for 30 min. Then 10 µL of fresh prepared iodoacetamide (IAA, 400 mM) solution was added and the mixture was incubated in the dark at room temperature for another 30 min. The sample was transferred to a 10kDa centrifuge filter, then we added 300 µL of 50 mM ammonium bicarbonate buffer to the filter and centrifuged at 14,000 × g for 10 min, repeated twice to remove excess salt. The samples were transferred from the filter to the tube, digested with 1 µg of sequencing grade trypsin (Promega V5111) and $CaCl_2$ (1 mM) at 37°C overnight. After heated at 95°C for 5 min, the digests were filtered through a 0.2 µm filter and vacuum-centrifuged to near dryness via a speed-vac vacuum concentrator. The samples were stored in -80°C for further analysis. The digests were re-dissolved in 20 µL Buffer A (99.9% $H_2O$, 0.1% formic acid) and loaded onto a Orbitrap Fusion mass-spectrometer. 3 µL of the sample was loaded onto a Thermo Acclaim PepMap100 precolumn (75 µM × 2 mm) and eluted on a Thermo Acclaim PepMap RSLC analytical column (75 µM × 15 cm). Buffer A (0.1% formic acid in $H_2O$) and Buffer B (0.1% formic acid in acetonitrile) were used to establish the 120 min gradient comprised of 90 min of 2−35% B, 8 min of 35−60% B, and 2 min of 60−95% B, 8 min of 95% B, followed by re-equilibrating at 2% B for 2 min. The flow rate was 0.3 µL/min. Peptides were than analyzed on a Thermo Orbitrap Fusion Lumos proteomic mass spectrometer in a data-dependent manner, with automatic switching between MS and MS/MS scans using a cycle time 3 s. MS spectra were acquired at a resolution of 240,000 with an AGC target value of $5\times10^5$ ions or a maximum integration time of 100 ms. The scan range was limited from 375 to 2000 m/z. Peptide fragmentation was firstly performed via a step higher-energy collisional dissociation (HCD) with the energy set at 15%, 30% and 45%, respectively, with an AGC target value of $5\times10^4$ and a maximum integration time of 48 ms. Additionally, collision induced dissociation (CID) fragmentation was triggered by detection of a 204.0865 ion signal, with the collision energy set at 35% and anactivation time of 10 ms. MS spectra were acquired at a resolution of 30,000 with an AGC target value of $5\times10^4$ ions and a maximum integration time of 54 ms. The raw data was analyzed using pGlyco 2.2.2 software, following the manual's instruction. In general, the protein database was from UniProt (ID P35613), the mass tolerance for precursors and fragment ions were set as ± 5 ppm. and ± 20 ppm., respectively. Maximal missed cleavage was set as 2. Carbamidomethylation on cysteine residues (C +57.022 Da) was used as a fixed modification, oxidation on methionine (M +15.995 Da) was a variable modification. The N-glycosylation sequon (N-X-S/T, X ≠ P) was modified by changing 'N' to 'J'. The N-glycosylation database was set as default in the software. The raw data and analysis files for proteomic analyses will be deposited to XXX.
